# Reticulate History of Southern South American Wild Potatoes: Recurrent Hybridization, Cytonuclear Discordance, and Phylogenetic Placement of *Solanum malmeanum*

**DOI:** 10.64898/2026.09.06.749666

**Authors:** Rodrigo Nicolao, Péter Poczai, Rocio Deanna, Madison Bullock, Morgan Gostel, Caroline M. Castro, Gustavo Heiden

**Affiliations:** Faculty of Agronomy Eliseu Maciel, Graduate Program in Agronomy, Federal University of Pelotas, P.O. Box 354, Pelotas, RS 96010-900, Brazil; Finnish Museum of Natural History & Department of Organismal and Evolutionary Biology, Faculty of Biological and Environmental Sciences, University of Helsinki, P.O. Box 7, FI-00014 Helsinki, Finland; Department of Biological Sciences, Texas Tech University, 2500 Broadway, Lubbock, TX 79409, USA; Morris Arboretum and Gardens, University of Pennsylvania, 100 E Northwestern Avenue, Philadelphia, PA 19118, USA; Botanical Research Institute of Texas, 1700 University Drive, Fort Worth, TX 76107, USA; Embrapa Clima Temperado, Rodovia BR-392, Km 78, 9° Distrito, Monte Bonito, P.O. Box 403, Pelotas, RS 96010-971, Brazil

**Keywords:** Angiosperms353, Chloroplast capture, Cytonuclear discordance, Introgression, Phylogenomic, Reticulation

## Abstract

Hybridization is a powerful process in plant evolution generating reticulate patterns of speciation, introgression, and chloroplast capture. In the *Petota* clade of *Solanum*, which spans cultivated and wild potatoes, these processes cause much of the taxonomic complexity of the group. We examined the evolutionary history of the *Commersoniana* group (*Solanum commersonii* and *S. malmeanum*) relative to *S. chacoense* (*Tuberosa* Group) using an integrative framework of phylogenomics and morphology. With target-capture sequencing (Angiosperms353) of nuclear and plastid loci recovered from herbarium specimens, together with data mining we built nuclear and plastome phylogenies and ran symmetry tests, gene concordance analyses, and coalescent simulations to weigh incomplete lineage sorting against chloroplast capture. The data reveal cytonuclear discordance and events consistent with chloroplast capture from *S. chacoense* into *Commersoniana* lineages supported by both empirical and simulated results. Phylogenetic networks confirm repeated historical and recent reticulation, especially involving *S. malmeanum*. Non-metric Multidimensional Scaling (NMDS) of morphological traits did not recover clear species boundaries but was consistent with a hybrid origin for the group. These results indicate that recurrent hybridization and event consistent with chloroplast capture are strongly influencing evolution in the examined wild potatoes, complicating classical taxonomy and informing conservation and breeding. They also underscore the value of combining genomic and phenotypic data to resolve species boundaries in plant groups with tangled hybrid histories.

## Introduction

*Solanum* section *Petota* (Solanaceae), now treated as *Solanum* clade *Petota* Dumort. [Bohs & Olmstead] under the PhyloCode **[1]**, comprises the cultivated potato (*S. tuberosum* L., Andigenum and Chilotanum Groups) and more than 100 wild species **[2, 3, 4, 5, 6]**, a vital reservoir of genetic diversity for the crop **[7, 8]**. Species counts have ranged between 112 and 232, depending on the author **[2, 3, 4, 5, 6]**. Correll recognized 157 wild species in 1962 **[2]**, Hawkes 217 wild plus 7 cultivated species across 21 series in 1990 **[3]**, Spooner and Hijmans 199 species in 2001 **[4]**, Spooner and Salas 188 species in 2006 **[5]**, and Spooner et al. 107 wild and 5 domesticated species in 2014 **[6]**. These species range across the Americas, from the southwestern United States (38° N) to southern Chile (41° S) and into Brazil, with the Andes from north-central Peru to central Bolivia as their center of diversity **[9]**.

Many potato species look alike yet stay partly isolated by pre- and post-zygotic barriers. Pre-zygotic barriers include pollen-pistil incompatibilities and ecogeographical isolation **[10, 11, 12]**. Post-zygotic barriers stem from abnormal endosperm development associated with Endosperm Balance Number (EBN)^1^ incompatibilities, but may also involve hybrid embryo abortion, reduced hybrid viability, and hybrid sterility **[6, 13]**. Potato evolution is influenced by intra- and interspecific hybridization, introgression, auto- and allopolyploidy, mixed sexual and asexual reproduction, and recent divergence **[17]**, which, together with rapid speciation across the heterogeneous Andean landscape, make the group exceptionally complex **[4]**. Hawkes **[3]** drew on Anderson’s “hybridization of the habitat” concept **[18]** and argued that hybrids persisting vegetatively through tubers and stolons increase the complexity. Many wild potatoes hybridize artificially into viable F1 and later generations **[19]**, yet subtle chromosomal differences and ecological specialization keep species apart **[20, 21]**.

Morphology provided the first evidence of natural hybridization in potatoes **[22]**, with Hawkes and Hjerting finding interspecific hybrids in 9.5% of specimens from Argentina, Brazil, Paraguay, and Uruguay **[20]**. Molecular work later confirmed that hybridization is common across *Petota* **[23, 24]**. Many species boundaries appear dubious and possibly arbitrary, inviting competing classifications **[2, 6, 20, 25]**, and this clade shows extensive reticulate evolution through both homoploid and polyploid hybridization **[3, 20, 21, 22, 23, 26, 27, 28, 29, 31]**.

Plant evolution is often reticulate **[32]**. Hybridization can spawn new species through novel trait combinations or polyploidization **[33, 34, 35]**, cause extinction by genetic swamping **[36]** or carry beneficial alleles between lineages **[37]**. It acts at both shallow **[38]** and deep **[39]** timescales and profoundly shapes diversification **[40, 41]**, fuelling genetic variation **[42, 43, 44]**, adaptation **[45, 46, 47]**, niche expansion **[48, 49]**, differentiation **[50]**, and speciation **[51, 52]**. Genomic advances have toppled the old assumption that speciation requires complete reproductive isolation, putting hybridization, introgression, and incomplete lineage sorting (ILS) at the heart of diversification **[53, 54]**.

Gottlieb proposed identifying hybrid taxa by sympatry with putative parents, intermediary morphology, partial F1 fertility, and biochemical additivity **[55]**. These signs, with phylogenetic incongruence among nuclear, chloroplast, and mitochondrial datasets, remain the main tools for detecting hybrids **[56, 57]**. Because plants carry three genomes whose histories may disagree **[58]**, cytonuclear discordance is common **[59]**. Potato evolution reflects a tangled interplay of ILS and interspecific hybridization **[2, 6, 21, 24, 26, 60, 61, 62, 63, 64, 65, 66]**, especially in polyploid origins **[67, 68, 69, 70, 71, 72, 73, 74, 75]**. Masuelli et al. argued that homoploid hybridization is the main factor responsible for the origin of diploid potato species **[76]**, a view later supported by Zhang et al. **[31]**. To date, 26 species (5 cultivated and 21 wild) are regarded as products of hybrid speciation **[6, 20, 25, 65, 77, 78, 79, 80]**. Even without leading to speciation, hybridization and introgression have been crucial to the adaptive success of potatoes across diverse environments **[21, 24, 65]**.

The Commersoniana group pairs two wild species, *Solanum commersonii* Poir. and *S. malmeanum* Bitter, across the Southern Cone of South America in Argentina, Brazil, Uruguay, and Paraguay **[6, 79]**. They occupy a broad variety of habitats, including *Araucaria angustifolia* (Bertol.) Kuntze forests and inundated savannas as well as grasslands, agricultural fields, roadsides, riverbanks, seashores, and coastal dunes, from sea level to 750 m, and produce flowers and fruits between October and July **[2, 20, 79]**. Plants are rosette to decumbent, with odd-pinnate leaves often bearing interstitial leaflets, long-acute to attenuate calyx lobes, white to lilac corollas (stellate in diploids, semi-stellate to pentagonal in triploids), globose to ovoid fruits, and 1 to 2 cm tubers at each stolon tip **[2, 3, 6, 20, 79]**. The status of *S. malmeanum* has shifted repeatedly, named as a distinct taxon **[81]**, *S. commersonii* f. *malmeanum* **[2, 82]**, and *S. commersonii* subsp. *malmeanum* **[20, 84]**, before being reinstated as *S. malmeanum* **[79]** (see **[84]** for review). Correll treated both as forms of *S. commersonii*, separating f. *malmeanum* by its at least partly petiolulate leaflets and often two or more interjected leaflets **[2]**. Hawkes and Hjerting kept them as subspecies, distinguishing *S. malmeanum* by leaflets that shrink gradually toward the base, narrowly decurrent to petiolulate, low to median pedicel articulation, a consistently white corolla, and geographical separation **[20]**. Mentz and Oliveira **[82]** noted that some glasshouse-grown *S. malmeanum* did not keep the white corolla, suggesting they might be hybrids of the two species, much as Correll had treated them **[2]**.

The wild species *S. chacoense* (2EBN, *Tuberosa* Group) grows in sympatry with *S. commersonii* (1EBN) and *S. malmeanum* (1EBN) yet stays reproductively isolated **[79, 85]**. Even so, the barriers within the group are leaky **[20, 86, 87]**, and *S. malmeanum* has been proposed to arise from natural hybridization between *S. commersonii* and *S. chacoense* **[2, 86]** (Fig. S7). All three taxa occasionally produce unreduced (2n) gametes that breach EBN-based barriers, yielding offspring that can remain fertile, introgress through backcrossing, or persist by vegetative propagation **[86, 87, 88]**. Such inter-EBN crosses likely shaped the history and diversity of these taxa, fostering adaptive traits in the wild and supplying valuable alleles for breeding into cultivated potato **[67, 68, 69, 70, 71, 72]**.

Morphologically, *S. commersonii* differs from *S. malmeanum* in its uppermost lateral leaflets that shrink rapidly toward the leaf base, its shorter lateral leaflets, and its purple corolla, whereas *S. malmeanum* has subequal uppermost leaflets that barely taper, usually petiolulate lateral leaflets, and white corollas. *S. chacoense* stands apart from *S. malmeanum* by its larger, erect habit, acute terminal-leaflet apex, and longer peduncles, and *S. commersonii* from *S. chacoense* by its smaller size, lateral leaflets that shrink rapidly toward the base, larger terminal leaflet, fewer lateral leaflets, and purple corolla **[79]**. The key morphological differences among *S. chacoense*, *S. commersonii*, and *S. malmeanum* are summarized in Table 1.

**Table 1.** Comparative morphological characters and geographical distribution of *Solanum commersonii*, *S. malmeanum*, and *S. chacoense* (Spooner & al. 2016). Countries: Argentina (ARG), Bolivia (BOL), Brazil (BRA), Paraguay (PAR), Peru (PER), Uruguay (URU).

| Character | *S. commersonii* | *S. malmeanum* | *S. chacoense* |
| --- | --- | --- | --- |
| **Habit / height** | Semi-rosette to erect, 0.15–0.3 m (to 1 m in shade) | Erect to semi-rosette, 0.15–0.3 m (to 1 m in shade) | Erect, larger, 0.5–2 m |
| **Uppermost lateral leaflets** | Decrease rapidly in size toward leaf base | Subequal, not decreasing rapidly toward base | Subequal except the most proximal 1–2 pairs |
| **Lateral leaflets** | Shorter (1–3.8 cm), mostly sessile | Larger (3.3–6.5 cm), generally petiolulate | 2.7–9 cm, petiolules 0–5 mm |
| **Lateral leaflet pairs** | 2–5 (fewer) | 3–5 | 4–7 |
| **Terminal leaflet** | Larger relative to laterals; obtuse apex | Obtuse (rarely acute) apex | Acute to acuminate apex |
| **Peduncle length** | 1.5–6.5 cm | 1.5–6 cm | 2.5–10.5 cm |
| **Corolla color** | Purple (occasionally white-tinged-purple) | White (rarely with a faint bluish spot) | White to creamy yellow |
| **Geographical Distribution** | ARG, BRA, URU | ARG, BRA, PAR, URU | ARG, BOL, BRA, PAR, PER, URU |

As these cases show, the taxonomy of *Solanum* clade *Petota* is notoriously snarled by hybridization, introgression, polyploidy, and morphological overlap, all of which have spawned competing classifications **[79]**. Untangling speciation in the Commersoniana group therefore demands an integrative approach that unites geographical, molecular, and morphological evidence **[2, 3, 6]**.

Plastid and nuclear phylogenomic analyses have consistently recovered *Solanum* section *Petota* as comprising several major lineages, including the distinct Clade 3 and Clade 4 South **[24, 31]**. However, the placement of species within these lineages remains unresolved. Zhang et al. **[31]** recovered *S. commersonii* and *S. chacoense* together in Clade 4 South, whereas *S. malmeanum* was placed in Clade 3. This unexpected phylogenetic position contrasts with the close morphological similarity between *S. malmeanum* and *S. commersonii*, their partially overlapping geographic distributions in the Southern Cone of South America, and previous hypotheses suggesting a close evolutionary relationship and possible hybrid origin involving these taxa **[2, 3, 6, 79, 81]**. Consequently, the phylogenetic position of *S. malmeanum* and the evolutionary relationships among these species remain uncertain, warranting an independent phylogenomic assessment.

Integrating modern phylogenomic tools, such as the Angiosperms353 target-capture bait set **[89, 90, 91],** and morphological analysis has elucidated taxonomic relationships, species boundaries, and the interpretation of morphological evolution across diverse groups of plant lineages **[92, 93]**. High-throughput sequencing approaches such as Hyb-Seq, particularly using target-probes for 353 low-copy nuclear genes (Angiosperms353), prove to be effective in generating informative data across any taxon of flowering plants [92]. Zhao & al. (2025) reconstructed ancestral states of morphological traits using their nuclear phylogeny. Their findings supported the hypothesis that large scale genomic changes and key morphological innovations may have contributed to adaptive evolution and increased biodiversity in rosids. Moreover, targeted enrichment of low-copy nuclear genes has been applied to elucidate potato species relationships and boundaries within the group at high resolution, clarify speciation, and deepen our grasp of the evolutionary history of wild potatoes **[91, 92, 93]**.

Here, we resolve the evolutionary relationships among the *Commersoniana* species (*S. commersonii* and *S. malmeanum*) and *S. chacoense* (Tuberosa Group) using an integrative framework combining phylogenomic data and morphology. We test whether the evolutionary history of *S. malmeanum* reflects hybrid ancestry involving *S. commersonii* and *S. chacoense*, or instead represents a distinct lineage shaped by historical and contemporary introgression. We further investigate whether the *Commersoniana* group represents a continuum of divergence rather than fully discrete lineages, and assess how hybridization, introgression, and incomplete lineage sorting have contributed to blurred species boundaries. We then evaluate whether *S. malmeanum* warrants recognition as a species distinct from *S. commersonii* and whether morphological characters traditionally used to distinguish taxa in *Petota* are congruent with phylogenomic evidence. By combining the Angiosperms353 target-capture platform with morphological analyses, we aim to clarify species boundaries and evolutionary processes in these wild potato species, with implications for their conservation and utilization in breeding.

## Materials and Methods

### Potato sampling

Reconstructing the evolutionary history of wild potatoes demands dense sampling across their native ranges, and thus, we drew on the extensive potato herbarium collection of the Wisconsin State Herbarium (WIS), complemented by a smaller number of specimens from the Field Museum of Natural History (F), Chicago, Illinois, and the Botanical Research Institute of Texas (BRIT), Fort Worth, Texas. Our strategy aimed to represent the native ranges of the ingroup taxa as broadly as possible. We sampled 13 *Solanum chacoense* specimens from the Tuberosa group, spanning latitudes from Peru to Argentina and reflecting the wide ecological range of the species. For the Commersoniana group, we sampled accessions of 5 *S. commersonii* and 11 *S. malmeanum*, respectively, which represent a comparable spread of ecological and geographic diversity, with several representatives from southern Brazil, Uruguay, and Argentina. We did not exhaustively cover every biogeographic region, but our sampling captured the areas known to harbour genetic and morphological variation in each species. We rooted the analyses with two outgroups, one representing Potato Clade 1+2 (*S. tarnii*) and the other Tomato Clade (*S. lycopersicum*). Many specimens were gathered by notable *Solanum* collectors^2^ whose well-documented field collections strengthen the taxonomic reliability of our dataset and tie it to earlier systematic work in the *Petota* clade.

To broaden taxon coverage, we added a small, select number of previously published RNA-seq data representing *Solanum* clade *Petota* clades 3 and 4 South, together with the outgroups *S. tarnii* (Clade 1+2) and *S. lycopersicum*, from Zhang et al. **[31]**. We downloaded the reads from the NCBI Sequence Read Archive (SRA) on the Puhti supercomputer, quality-filtered them with FastP **[94]**, and assembled them with HybPiper v2.3.2 **[89]** to recover nuclear loci from the Angiosperms353 set and plastid coding sequences. We aligned the recovered sequences with MACSE under a codon aware framework using the OMM (Ongoing Multiple Alignment Model) **[95]**. To screen for violations of model assumptions, we concatenated the gene alignments with BAD2matrix **[96]**, ran symmetry tests in IQ-TREE v2.3.6, and kept only the genes that passed for subsequent downstream phylogenetic analyses.

The geographic origin of each accession is shown in Fig. 1 using the codes from Table S1.

**Fig. 1.**
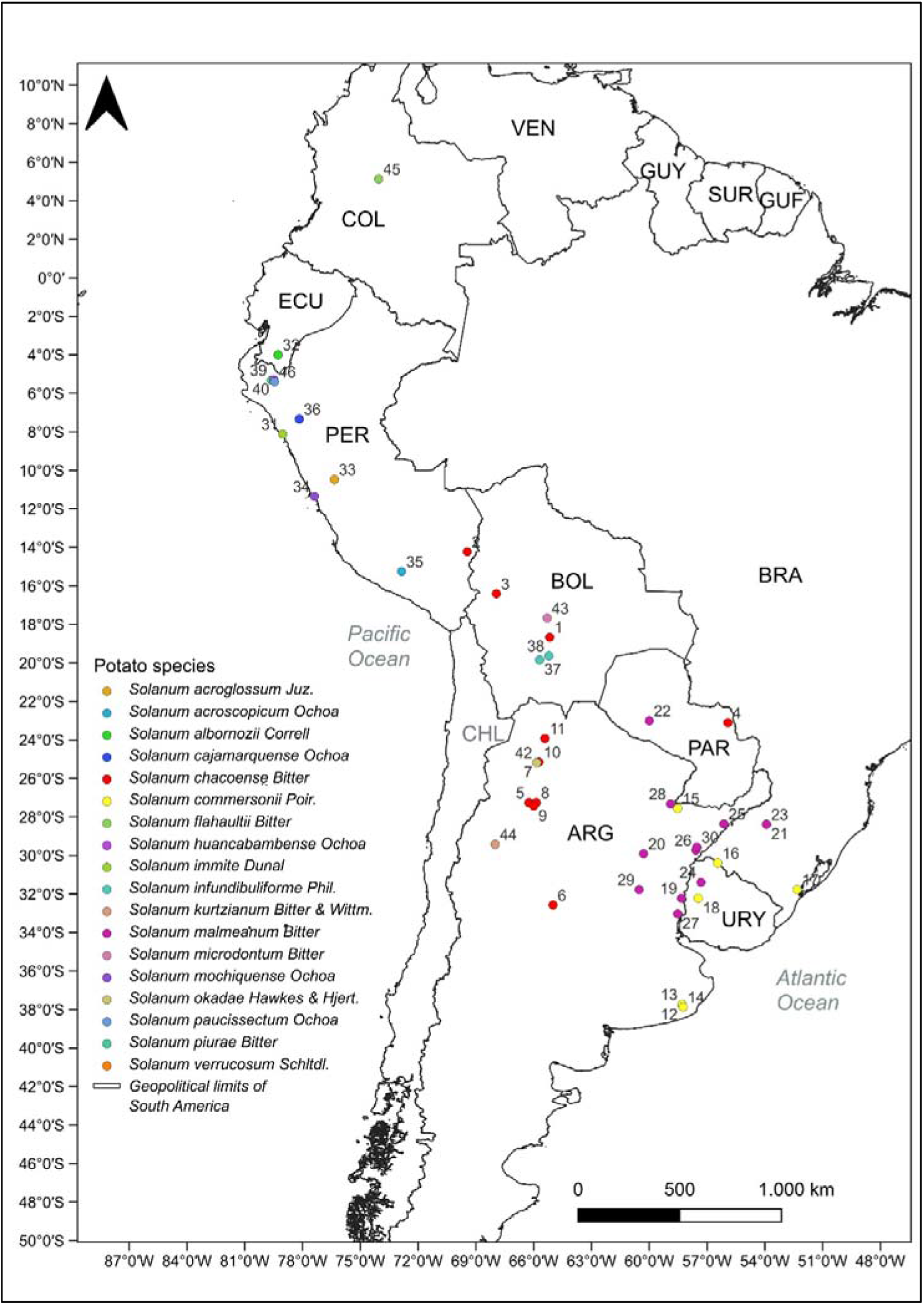
Geographic distribution of sampled wild *Solanum* species across South America. Colored circles indicate collection localities of accessions, labeled by species code (sample number; see Table 1). Colors correspond to potato species as indicated in the legend. Country abbreviations are as follows: PER (Peru), BOL (Bolivia), ARG (Argentina), CHL (Chile), PAR (Paraguay), URY (Uruguay), BRA (Brazil), ECU (Ecuador), COL (Colombia), VEN (Venezuela), GUY (Guyana), SUR (Suriname), and GUF (French Guiana). Geopolitical boundaries are shown in black lines. Species names correspond to the accession codes listed in Table 1.

### DNA extraction, library preparation, and sequencing

We extracted total genomic DNA from herbarium leaf material with the Qiagen DNeasy Plant Mini Kit (Hilden, Germany) at the George C. and Sue W. Sumner Molecular and Structural Laboratory of the Botanical Research Institute of Texas (BRIT), following the manufacturer protocols. DNA concentration was measured using a Qubit fluorometer and the dsDNA HS Assay from Life Technologies (Grand Island, NY, USA), also at the Sumner Laboratory at BRIT. Library preparation and Angiosperms353 (A353) target-capture sequencing **[90]** were carried out at Texas Tech University (TTU), where we used multiplexing **[97]** to reduce costs per sample.

### Read quality control and decontamination

Clean input is critical for herbarium data, and thus, we ran a multi-step decontamination protocol **[98]**. We first screened reads with FastQ Screen v0.16 **[96]**, which maps short reads against a panel of reference genomes with Bowtie2 or BWA to flag contamination. Our reference panel covered common laboratory and environmental contaminants, including *Homo sapiens*, *Mus musculus*, *Rattus norvegicus*, *Escherichia coli*, *Saccharomyces cerevisiae*, several rRNA databases, PhiX, vector sequences, and sequencing adapters (Decontamination 1). The tabular and graphical outputs let us quantify the proportion of reads mapping uniquely, multiple times, or not at all to each reference, and we flagged libraries with high proportions of ambiguous or non-target hits for further filtering. A second round of screening used the filtered outputs from the first step (Decontamination 2; Fig. 2) to consistently strip the herbarium-specific contaminants identified by Bieker et al. **[99]**. We then trimmed adapters, removed low-quality reads, and eliminated known technical artifacts with BBduk from the BBTools pipeline **[101]**, sharpening the accuracy of the downstream alignment and assembly of the Angiosperms353 data.

**Fig. 2.**
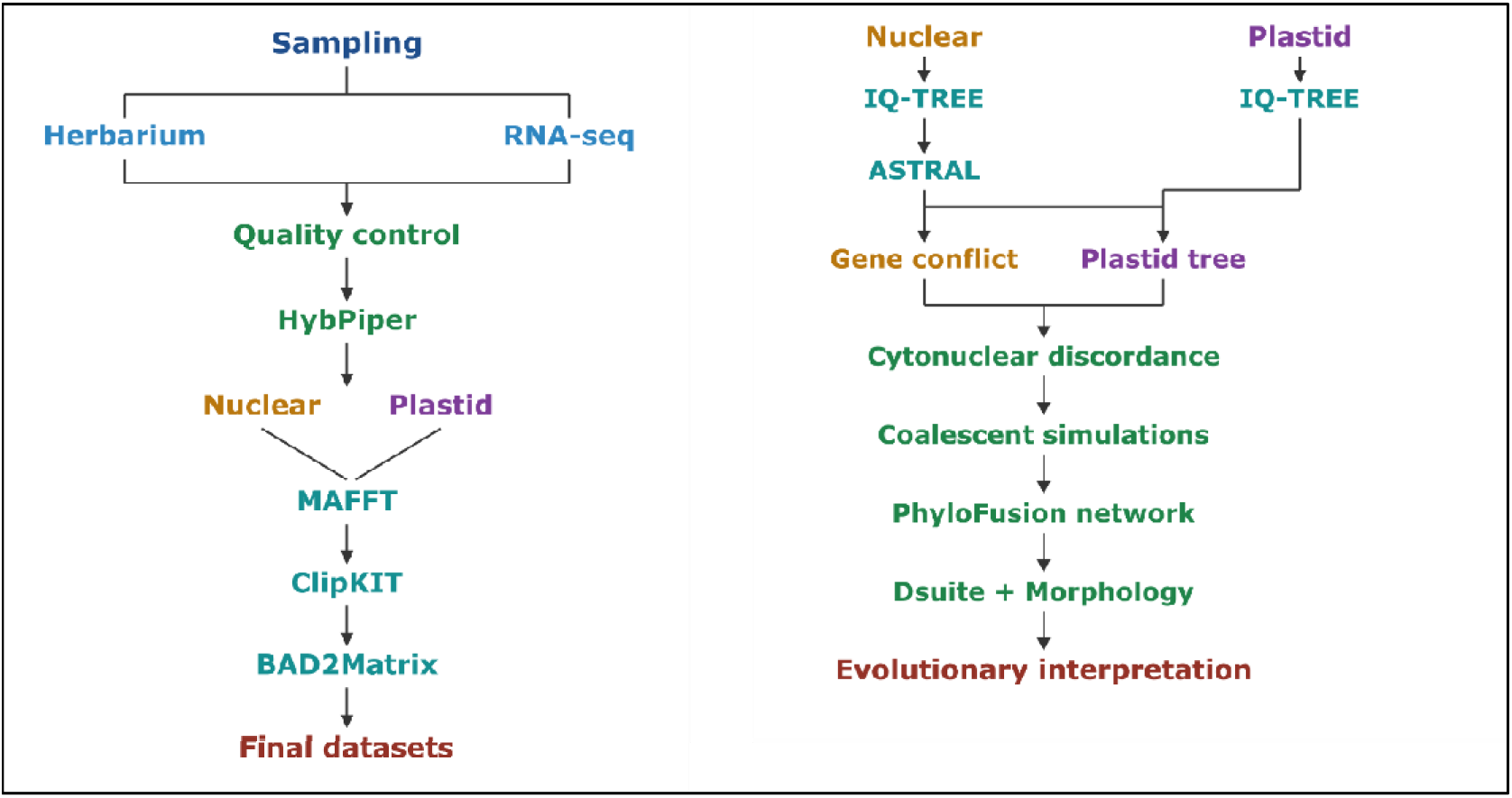
Bioinformatic workflow for phylogenomic analyses, from sampling and quality control through target-capture assembly (HybPiper), alignment (MAFFT), trimming (ClipKIT), and matrix concatenation (BAD2Matrix), generating the final nuclear and plastid datasets. Nuclear and plastid gene tree inference (IQ-TREE), coalescent-based species tree estimation (ASTRAL) for the nuclear dataset, and assessment of cytonuclear discordance through coalescent simulations, network inference (PhyloFusion), and introgression tests (Dsuite) combined with morphological data, culminating in the evolutionary interpretation of the results. Scheme generated in BioRender.

### Assembly and alignment of target-capture data

We recovered target genes with HybPiper v2.3.2 **[89]**, which maps reads to references with BWA **[102]** and assembles each locus *de novo* with SPAdes **[103]**, yielding exon and supercontig sequences that include introns and flanking regions wherever coverage allows. We ran HybPiper separately for the nuclear and plastid regions, recovering the latter by mining off target reads that mapped to the chloroplast genome **[104]**, and decreased the default coverage cutoff to 5 to maximize the number of retained contigs **[105, 106]**. We took the Angiosperms353 target file, assembled for tomato (*Solanum lycopersicum*), from the Royal Botanic Gardens, Kew Tree of Life Explorer **[107]**, and built the plastome target file from all protein coding regions of the tomato reference genome (RefSeq NC_007898.3). We aligned the recovered sequences for each gene with MAFFT v7.505 **[108]** under the *--auto strategy*, then trimmed poorly aligned and uninformative sites with ClipKIT v1.3.0 **[109]** under the smart-gap setting while preserving signal-rich regions. Finally, we compiled the per locus alignments into concatenated matrices and partition files with BAD2Matrix **[96]** for input to IQ-TREE. All analyses ran on the Puhti supercomputer at CSC in Espoo, Finland.

### Phylogenomic analyses

We analysed both the exon and supercontig assemblies of the nuclear and plastid datasets. For each plastid dataset, we inferred a concatenated maximum likelihood tree in IQ-TREE v2.3.6 **[110]** from 10 runs under the best fit model chosen per gene by ModelFinder **[111]** and assessed support with 1000 ultrafast bootstrap replicates **[112]** under a nearest neighbour interchange optimization. For the nuclear dataset, we estimated a species tree under the multispecies coalescent (MSC) in ASTRAL v5.7.3 **[113]**. We built individual gene trees for the recovered A353 loci in IQ-TREE with 1000 bootstrap replicates, conservatively dropping every gene that HybPiper flagged with a paralog warning, and summarized them into a species tree using the comprehensive annotation (-t 2) option of ASTRAL.

### Assessment of concordance and conflict

To probe the processes shaping wild potato evolution, we quantified the agreement and disagreement between individual gene trees and the species tree since such conflict can expose the footprints of incomplete lineage sorting, hybridization, or gene tree estimation error. We measured nuclear gene tree concordance and conflict with PhyParts **[114]**, applying a minimum bootstrap threshold of 50% (-s 0.5) to discard poorly supported branches and retaining only bipartitions present in at least one gene tree. We visualized the output with the *phypartspiecharts.py* script **[115]**, which draws on Matplotlib **[116]** and ETE3 **[117]** to place a pie chart at each internal node summarizing the proportion of gene trees that are concordant, conflicting, or uninformative. To quantify branch support at both the site and gene levels, we computed site concordance factors (sCF), gene concordance factors (gCF), and bootstrap support in IQ-TREE v2.3.6 **[118]**. The three metrics were then visualized together using the *concordance.R* script published by Lanfear **[119]** for a branch-wise comparison of support across the tree.

### Chloroplast tree simulations

To uncover why the plastid and nuclear phylogenies disagree, we paired concordance factor analysis with coalescent simulations. Following the framework of Folk et al. **[120]**, we asked whether incomplete lineage sorting (ILS) alone could explain the discordance between the plastid and nuclear trees or whether chloroplast capture was also required. We simulated plastid gene trees under the MSC for 10 000 generations with the contained_coalescent_tree function of DendroPy v5.0.8 **[121]**, using the nuclear ASTRAL species tree as a reference and lengthening its branches fourfold to reflect the smaller effective population size (N_e_) of plastid genomes. Because plastomes are haploid and typically maternally inherited, they carry roughly one quarter of the N_e_ of biparentally inherited diploid nuclear genomes, and since coalescent time scales inversely with N_e_, lineage sorting runs about four times faster in plastomes than in nuclear loci **[122]**. Scaling the species tree this way lets the simulated plastome trees mirror plastid coalescent dynamics. We then computed the Robinson-Foulds (RF) distance **[123]** between each simulated plastome tree and the nuclear tree to build a null distribution of topological distances expected under ILS and compared the empirical plastome to nuclear RF distance against it. An observed distance that significantly exceeds the null indicates that ILS alone cannot account for the discordance, pointing instead to plastome capture or hybridization.

In parallel, we quantified how often each branch of the empirical plastome tree appeared among the simulated gene trees with the -gcf function of IQ-TREE v2.3.6, giving a branch-wise concordance analysis. Under the null hypothesis that ILS alone generates the discordance, conflicting branches in the plastome tree should recur frequently among the simulations. Branches recovered at low frequency (≤1%) therefore led us to reject the null hypothesis and to infer that additional processes, such as chloroplast capture, contributed to the cytonuclear discordance **[124]**.

### Network analysis

To test for hybridization directly, we fused the nuclear and plastid phylogenies with PhyloFusion **[125]** as implemented in SplitsTree v6.4.17 **[126]**. We retained a single network under the mutual refinement criterion, using a refined heuristic search that groups non-separated taxa and applies clade reduction, which yields a directed acyclic graph whose arcs represent hybridization events. Onto this network, we overlaid the conflicting branches recovered at low frequency (≤1%) in the plastome simulations to highlight the events attributable to chloroplast capture.

### Detecting introgression in the nuclear genome

To test whether the chloroplast-nuclear discordance recovered by the network reflects introgression in the nuclear genome rather than incomplete lineage sorting, we built a nuclear SNP matrix and screened it for admixture. The A353 reads were first screened against the tomato chloroplast and mitochondrial genomes (mitochondrial accession MF034193) with BBMap **[127]** to retain the nuclear fraction, then aligned to the *S. commersonii* reference genome (NCBI assembly ASM1825827v1) with Bowtie2 **[128]** in local mode under the very-fast-local preset (match bonus 2, mismatch penalty 6, ambiguous-base penalty 1, gap open and extend penalties 5 and 3, Phred+33, unmapped reads discarded). Alignments were sorted with SAMtools **[129]**, rescaled for DNA damage with mapDamage 2.0 **[130]**, and reduced to reads with a mapping quality of at least 20 after removing secondary alignments. Variants were called with BCFtools **[129]** using mpileup (maximum depth 100, minimum base and mapping quality 20) and the diploid multiallelic caller restricted to SNPs, then filtered to biallelic sites with a total depth of at least 5, genotypes with a PL-derived quality difference above 10, and a minor allele frequency above 0.07 in VCFtools **[131]**.

We tested for introgression with Dsuite **[132]**, using the nuclear ASTRAL tree as guide topology. Dtrios computed Patterson’s D and the f4-ratio for all trios across *S. commersonii*, *S. malmeanum*, and *S. chacoense*, with the hybrid accessions *S. commersonii* (Boldrini & Boechat 264) and *S. malmeanum* (Delhey s.n.) placed in their own groups so that each was compared against the remaining members of its species, and significance was taken from a block-jackknife Z-score over 20 blocks. Dinvestigate then localized the two hybrid signals in sliding windows of 50 SNPs stepped by 25, reporting f*d, f*dM, and d*f*.

### Morphological analysis, ordination plots, and taxon description

We selected 44 morphological traits, both continuous and discrete, that have featured in traditional taxonomic treatments of the *Solanum* clade *Petota*, comprising 30 vegetative and 17 reproductive characters **[79, 133]** (**Table S2**). We scored character states from herbarium specimens of three wild potato species, *S. commersonii* and *S. malmeanum* (Commersoniana) and *S. chacoense* (Tuberosa), measuring continuous traits with a digital INSIZE caliper (model 1108-150). The final matrix comprised 8 discrete and 41 continuous traits, five of the latter expressed as ratios.

To assess morphological differentiation among the wild potato populations, we ran non-metric multidimensional scaling (NMDS) on the subset of continuous traits with strong phylogenetic signal (**Suppl. Method 1**, and **Table S2**) together with all discrete traits (**Tables S3 and S4**). We standardized the continuous traits beforehand and computed morphological distances with the Gower metric, which handles mixed variable types **[142]**. We ordinated with the metaMDS function of the vegan package **[143]** and inspected stress values to decide how many dimensions adequately represented the data. To link individual traits to population differentiation, we fitted each trait with the permutation-based envfit function and treated traits with *p < 0.05* as significantly associated with the NMDS axes. We visualized the ordinations with ggplot2 **[144]** and ran all analyses in RStudio v2025.5.0 **[139]**. We characterized morphological variation among the studied populations with the MorphoTools2 package **[145]** in RStudio v2025.5.0 **[139]**.

## Results

### Data assembly

We generated target-capture data for 30 samples spanning four wild potato species of the Commersoniana group, the Tuberosa group, and the outgroup. Recovery averaged 309 genes per sample (246 to 347) and 200 708 base pairs (27 160 to 532 188); per-sample statistics for nuclear and chloroplast genes appear in **Table S1** and **Suppl. Fig. 1** and **Table S2** and **Suppl. Fig. 2**, respectively. After removing paralogous loci with HybPiper scripts **[89] (Suppl. Fig. 3 and Suppl. Fig. 4)**, the final dataset held 347 nuclear exon and 254 nuclear supercontig loci, plus 74 plastid exon and 62 plastid supercontig loci.

### Alignment statistics with IQ-TREE

Alignment statistics for the nuclear (A353) and chloroplast datasets, comparing exons with supercontigs, are summarized in **Table S3**.

#### Nuclear sequence alignment (A353)

The A353 supercontig alignment spanned 943 642 bp, mostly conserved (84% constant) yet retaining 11 219 parsimony-informative sites and a Townsend informativeness score of 0.69, a good basis for resolving relationships at intermediate depths. The exon alignment was shorter (209 286 bp) and more conserved (94.4% constant), with less missing data (17.6%) but only 345 informative sites and a slightly lower Townsend informativeness score (0.57). In IQ-TREE, the supercontig alignment clearly outperformed the exons, yielding higher mean bootstrap (93.6) and SH-aLRT (98.7) and a well-resolved, internally consistent tree, whereas the exon tree performed acceptably (bootstrap 80.6, SH-aLRT 89.96) but appeared to miss some deeper signal (**Suppl. Fig. 5)**.

#### Chloroplast sequence alignment

The plastid datasets showed the same pattern. The supercontig alignment (105 014 bp) was largely conserved (91.7% constant) but kept 8672 variable and 710 parsimony-informative sites, with a high Townsend informativeness score (0.718) despite substantial missing data (40.4%). The protein-coding alignment was far shorter (63 562 bp) and more conserved (97.4% constant), with only 39 of 1651 variable sites parsimony-informative, although it carried fewer missing data (26.4%). Its ML tree (recovered using IQ-TREE) reached acceptable bootstrap support (83.2) but an uneven SH-aLRT mean (69.3, SD 39.5), reflecting weak variation in some clades, while the supercontig tree gave similar bootstrap (84.7) and better SH-aLRT (77.5) (**Suppl. Fig. 5**).

### Nuclear species tree

The A353 supercontig species tree provided the backbone phylogeny for the Commersoniana group (*S. commersonii* and *S. malmeanum*) and *S. chacoense* (Tuberosa group), with *S. stoloniferum* (Longipedicelata group) as outgroup, against which the plastid relationships were compared. The Commersoniana group was monophyletic, whereas *S. chacoense* was paraphyletic (**Fig. 3**).

**Fig. 3.**
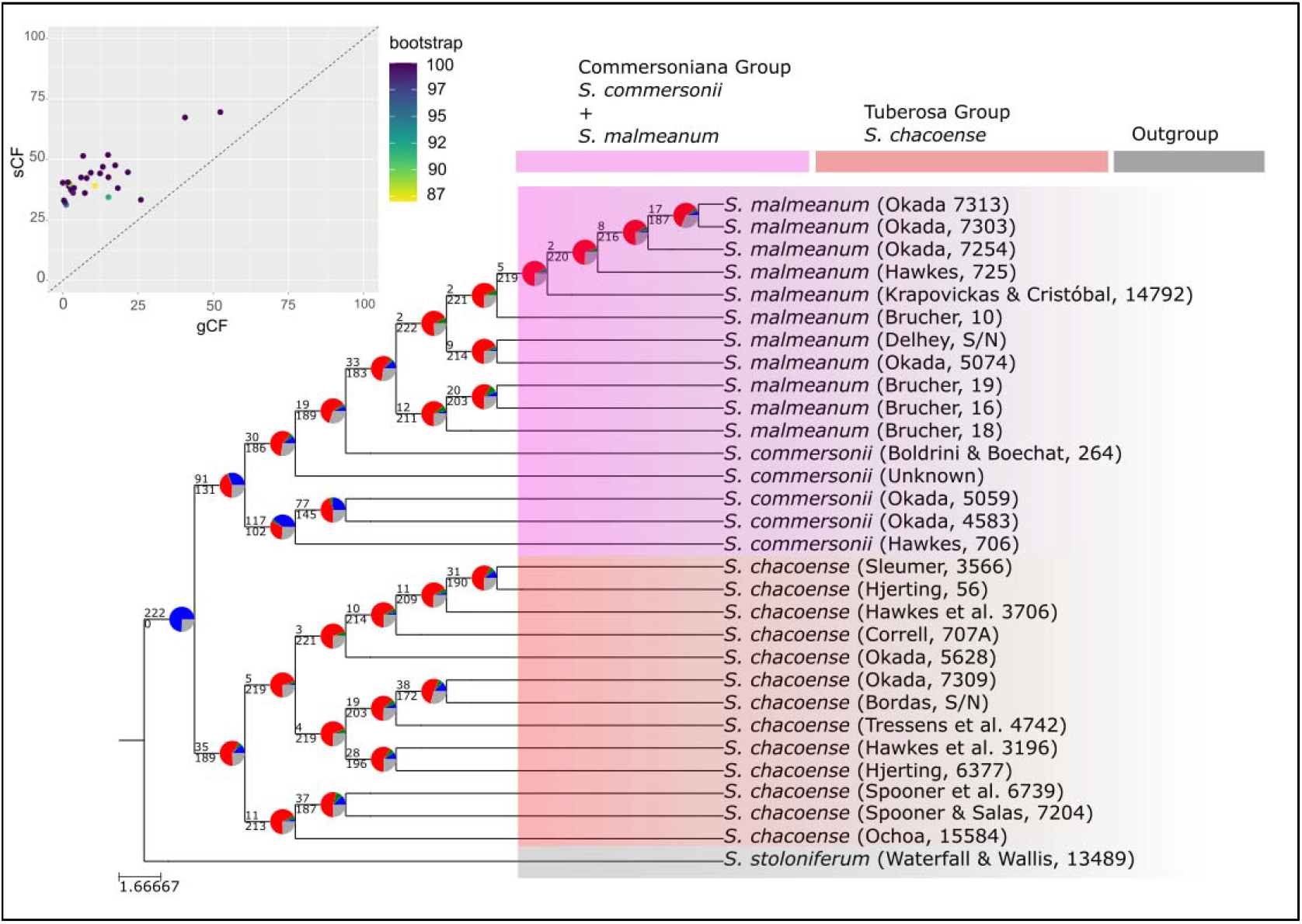
Nuclear species tree of the wild potato species, with the ingroups Commersoniana group (*S. commersonii* plus *S. malmeanum*) and *S. chacoense* (Tuberosa group) and the outgroup *S. stoloniferum* (Longipedicelata group) and *S. lycopersicum* (Tomato clade). Pie charts at each node give the proportion of gene trees concordant with the species tree (blue), supporting the most prevalent alternative bipartition (green), supporting all other discordant bipartitions combined (red), and uninformative for that node (gray). Numbers above and below branches are the counts of concordant and conflicting gene trees, respectively. The upper-left scatterplot compares site (sCF) and gene (gCF) concordance factors, each point a node positioned by sCF (y axis) and gCF (x axis) and shaded by bootstrap support. Nodes with high bootstrap but low concordance reflect gene tree conflict, likely from incomplete lineage sorting or hybridization, and help interpret plastid-nuclear discordance where the plastid signal departs from nuclear expectations.

To assess support beyond the bootstrap, we calculated site (sCF) and gene (gCF) concordance factors, which give the percentage of genomic sites and gene trees that support each species-tree branch rather than a conflicting alternative; the node pie charts show gene trees concordant with the species tree (blue), supporting the top alternative (green), supporting all other alternatives combined (red), and uninformative for that node (gray) (**Fig. 3**). Support formed a mosaic of strong and weak splits. Most nodes carried very low gene concordance but higher site concordance, and in the sCF against gCF plot the majority fell above the diagonal, where site support exceeds gene support **[116]**. This pairing of low gCF with high sCF means that branches retain strong site-wise signal even where individual gene trees conflict, a signature of reticulate evolution typical of groups shaped by hybridization, introgression, or ILS **[159]**.

The split between *S. chacoense* and the Commersoniana group was strongly concordant, its pie charts dominated by blue, whereas deeper nodes within each group were far more conflicted and dominated by red (**Fig. 3**). Discordant gene trees outnumbered concordant ones at the early branching *S. chacoense* node (189 versus 31), within the Commersoniana group (131 versus 91), and at the sister node of the *S. malmeanum* clade (183 versus 33), while the *S. commersonii* node was only marginally concordant (112 versus 102). Two *S. commersonii* samples (CST s.n. and Boldrini & Boechat 264) shared nuclear ancestry with *S. malmeanum*, their sister nodes likewise dominated by discordant gene trees.

Although bootstrap support was uniformly high across the species tree, the concordance factors varied widely, and several strongly supported nodes (bootstrap 87% to 100%) carried low gCF and sCF **(Suppl. Fig. 6) [159]**.

### Placement within the broader potato phylogeny

To place our focal species within the broader potato phylogeny, we recovered the A353 exons from the transcriptomic reads of Zhang et al. **[31]** and added these accessions to our target-capture matrix before reinferring the tree. The combined analysis recovered the two deep groups reported by Zhang et al. **[31]**, Clade 4 South and Clade 3, with *S. tarnii* (potato Clade 1+2) and *Solanum lycopersicum* (tomato) as successive outgroups **(Fig. 4)**. Within Clade 4 South, our data again recovered a well-supported, monophyletic Commersoniana group of *S. commersonii* and *S. malmeanum*, nested alongside a grade comprising *S. chacoense* and other Clade 4 species such as *S. boliviense*, *S. flahaultii*, *S. verrucosum*, *S. okadae*, *S. kurtzianum*, and *S. microdontum*. This contrasts with Zhang et al. **[31]**, who recovered *S. commersonii* in Clade 4 and *S. malmeanum* in Clade 3 as members of separate clades rather than a single lineage.

**Fig. 4.**
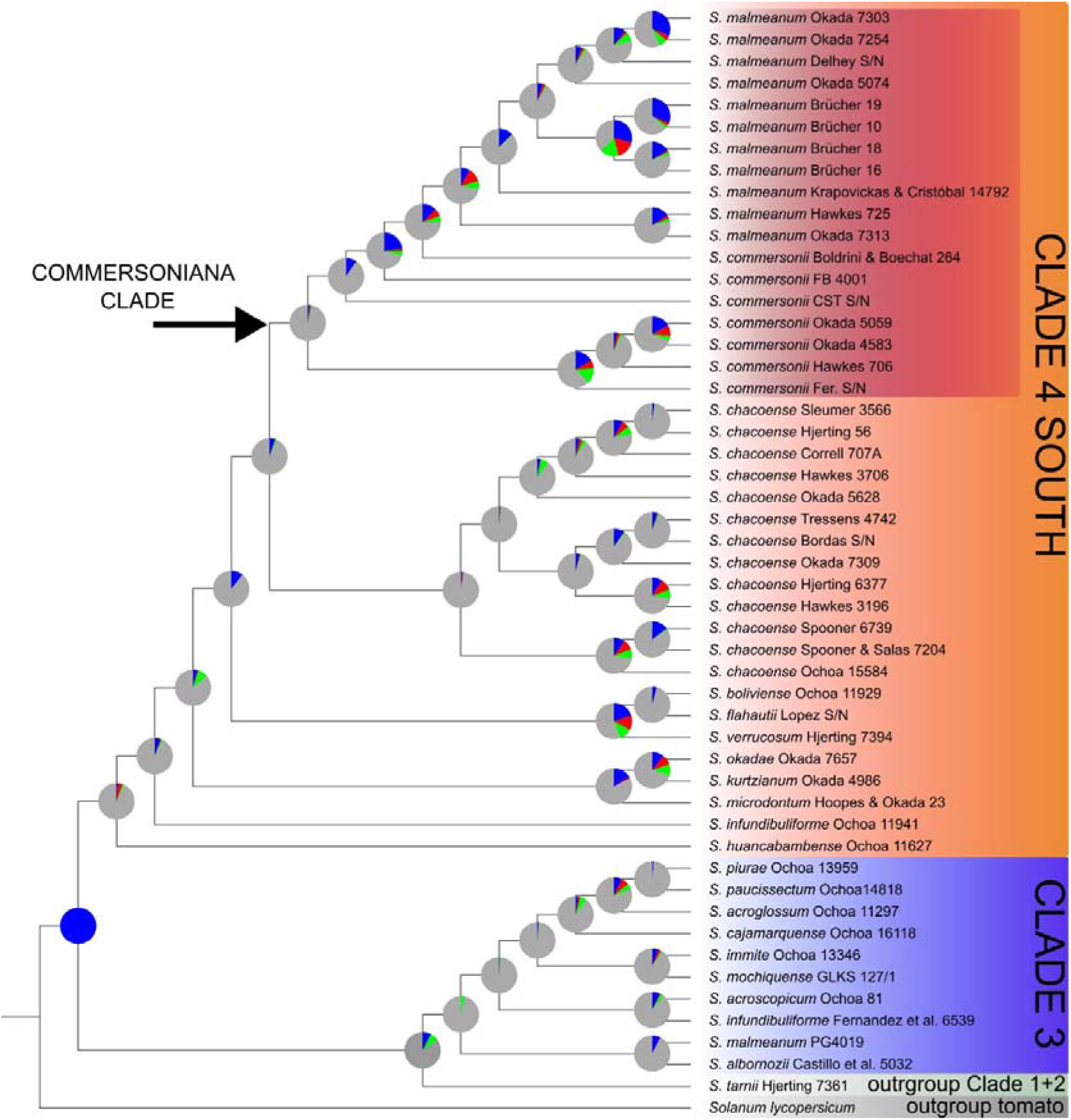
Nuclear species tree of wild potato species based on A353 nuclear loci recovered from target-capture and transcriptomic datasets. The phylogenomic analysis includes the focal Commersoniana group (*Solanum commersonii* and *S. malmeanum*), *S. chacoense* (Tuberosa group), and additional representatives of potato Clade 4 South and Clade 3. *S. tarnii* (potato Clade 1+2) and *S. lycopersicum* (Tomato clade) were included as successive outgroups. Pie charts at each node indicate the proportion of gene trees supporting the species-tree topology (blue), the most frequent alternative bipartition (green), all other conflicting bipartitions combined (red), and uninformative gene trees (gray). Numbers above and below branches represent the numbers of concordant and conflicting gene trees, respectively. The inset scatterplot compares site concordance factors (sCF) and gene concordance factors (gCF), with each point representing a node positioned according to its sCF (y-axis) and gCF (x-axis) values and shaded according to bootstrap support.

The only *S. malmeanum* accession to fall outside the Commersoniana group in our combined tree was PG4019, taken from the transcriptomic dataset, which was placed within Clade 3 far from every other *S. malmeanum* **(Fig. 4)**. Because our target-capture data for the same accession (Okada 5074) grouped with the remaining *S. malmeanum* inside the Commersoniana clade, this isolated placement most likely reflects a sample mix-up or mislabeling, e.g. during greenhouse propagation, rather than a genuine phylogenetic signal. The disagreement between the two studies over the position of *S. malmeanum* is therefore attributable to this single misplaced accession, and our denser sampling supports the monophyly of the Commersoniana group.

### Chloroplast phylogeny

In the chloroplast tree, the *S. chacoense* clade (Tuberosa group) was polyphyletic and split into two well-supported, except for Ochoa 15584, geographically structured groups, hinting at population differentiation, local adaptation, or ongoing speciation. A northern clade of two samples (Spooner & Salas 7204 and Spooner 6739) is fully supported (sCF = 100, bootstrap = 100), a southern clade of five samples (Correll 707A, Hawkes 3706, Hjerting 56, Sleumer 3566, and Okada 5628) is also strongly supported (sCF = 100, bootstrap = 99), and a third clade of four samples (Bordas s.n., Hawkes 3196, Hjerting 6377, and Okada 7309) is again fully supported (sCF = 100, bootstrap = 100).

The Commersoniana group likewise forms a fully supported clade (sCF = 100, bootstrap = 100) that divides into two subclades comprising accessions of *S. malmeanum* (sCF = 88, bootstrap = 82) and *S. commersonii* (sCF = 99, bootstrap = 92), respectively. One exception stands out, the *S. malmeanum* accession Delhey s.n., whose plastome groups with *S. commersonii*.

### Simulated chloroplast trees

To test whether the chloroplast-nuclear discordance could reflect incomplete lineage sorting (ILS) alone, we simulated chloroplast genealogies under a multispecies coalescent and compared their Robinson-Foulds (RF) distances to the species tree against the observed value (**Fig. 6**). Across 10 000 simulations, the mean RF distance was 58.140 ± 1.280, whereas the empirical chloroplast tree sat at 36.000, and no simulated tree fell below the observed value (p = 0.001). The empirical plastid tree is therefore far more congruent with the nuclear tree than ILS predicts, rejecting ILS as the sole cause of the cytonuclear discordance **[160]**. This high measure of congruence instead suggests chloroplast capture, in which historical introgressive hybridization replaces one species’ plastome with another’s **[161]** and pulls the plastid topology toward the nuclear species tree **[162]**, as seen here.

**Fig. 5.**
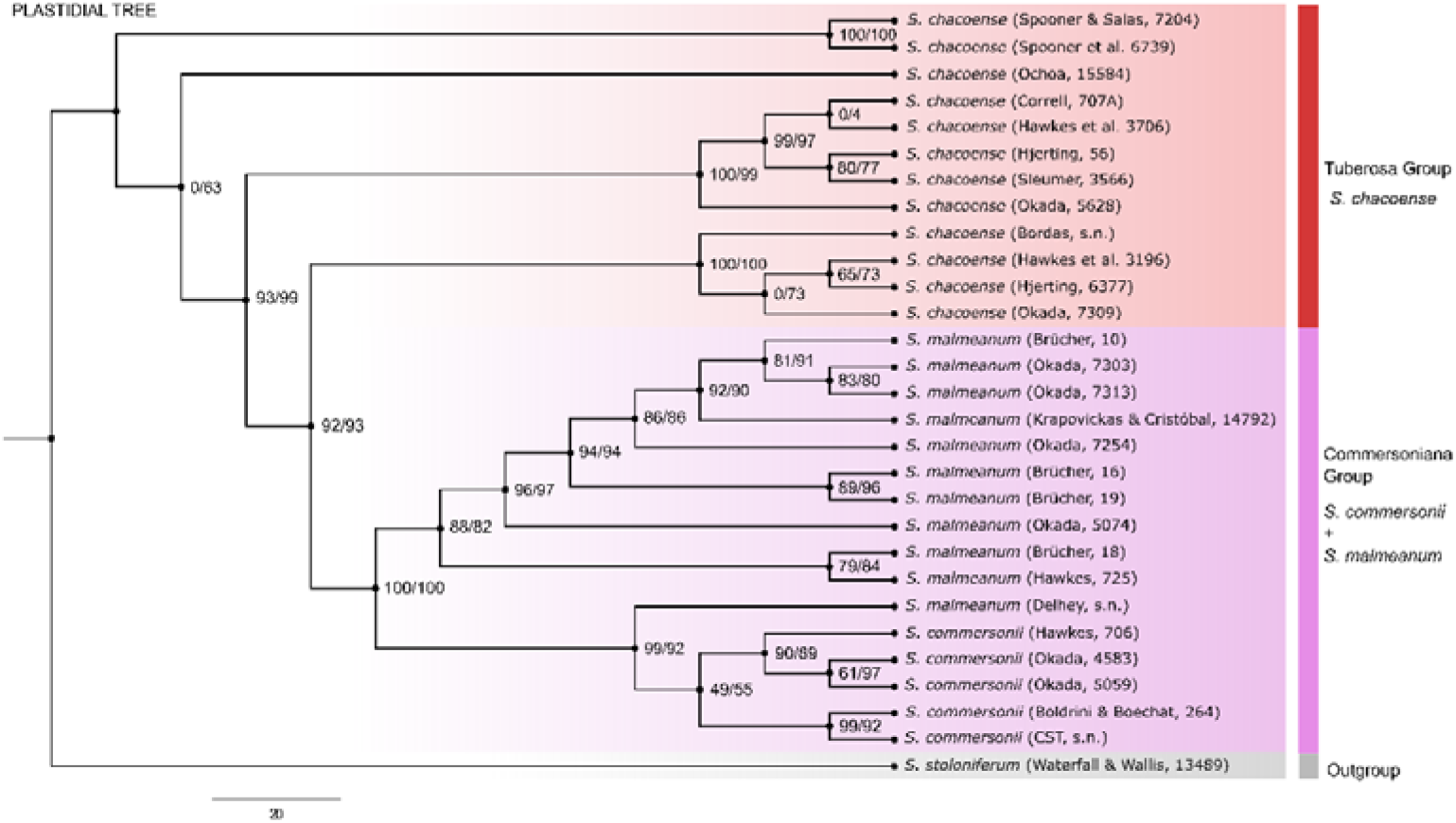
Plastidial phylogeny of the studied wild potato species, with the ingroups Commersoniana group (*S. commersonii* plus *S. malmeanum*) and *S. chacoense* (Tuberosa group) and the outgroup *S. stoloniferum*. Each node shows sCF and bootstrap support. The Tuberosa group (red) comprises *S. chacoense* in three main clades (chc 1, chc 2, chc 3), the Commersoniana group (pink) comprises *S. malmeanum* (mlm) and *S. commersonii* (cmm), and the outgroup (gray) is *S. stoloniferum*. The *S. chacoense* accession Tressens 4742 was excluded from the chloroplast and PhyloFusion trees because it recovered no plastid data.

**Fig. 6.**
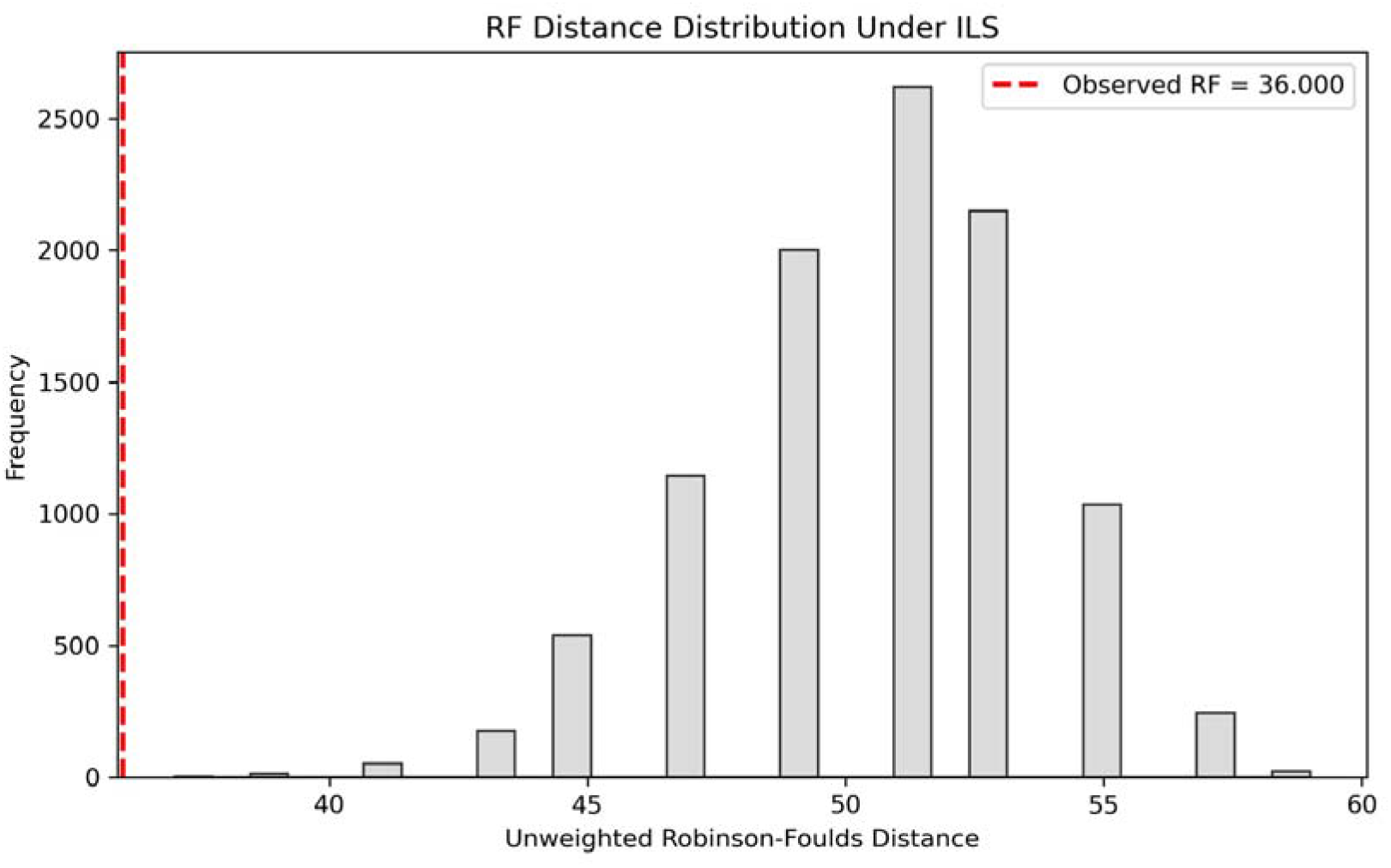
Coalescent simulation of chloroplast genealogies under a reduced plastid effective population size. Histograms show the distribution of Robinson-Foulds (RF) distances between the nuclear species tree and 10 000 simulated chloroplast gene trees, with the observed empirical RF marked by a red dashed line. Under ILS alone, the observed value should fall within the simulated distribution. Instead, it lies below nearly all simulated values, especially under the 4x scaling shown here (plastid Ne one-quarter of nuclear), so the empirical chloroplast tree is more congruent with the species tree than ILS predicts, pointing to additional processes such as introgression and chloroplast capture.

### Phylogenetic network inference with PhyloFusion

Fusing the nuclear and chloroplast phylogenies in PhyloFusion exposed extensive reticulation among these wild potatoes and pervasive conflict across the Commersoniana and Tuberosa clades. The network resolved 22 putative chloroplast capture events (red edges) and 4 ILS events (blue edges), each marking chloroplast-nuclear discordance left by historical hybridization (**Fig. 7**).

**Fig. 7.**
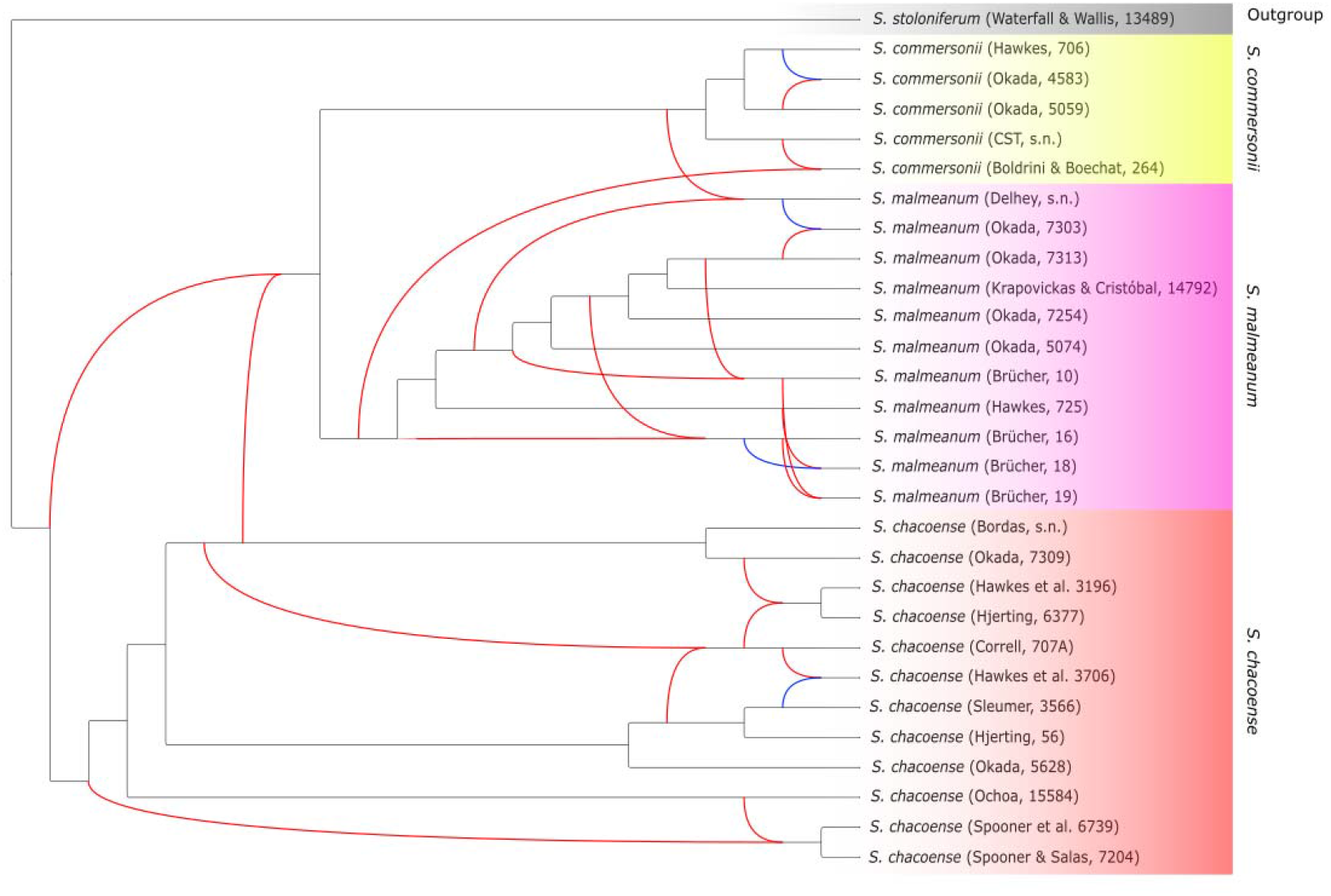
Phylogenetic network of the Southern South American wild potato species from chloroplast and nuclear data. The PhyloFusion of the chloroplast maximum-likelihood tree and the nuclear species tree reveals multiple reticulations, 22 hybridization events (red edges), and four ILS events (blue edges) across the species.

Two *S. commersonii* accessions (CST s.n. and Boldrini & Boechat 264) sit inside the *S. malmeanum* clade based on nuclear data, yet their plastomes place them firmly within the clade that comprises *S. commersonii* (sCF = 99, bootstrap = 92). Another example is *S. malmeanum* Delhey s.n., which carries a *S. commersonii*-type plastome despite a clear *S. malmeanum* nuclear assignment. These opposing mismatches indicate independent reticulations consistent with chloroplast capture events, with plastid introgression across species boundaries through hybridization and backcrossing while the nuclear genomes stayed distinct. Where both concordance factors were low and the corresponding plastid relationship was almost never recovered in our coalescent simulations (1% or less), chloroplast capture is a more plausible explanation than stochastic ILS. Concordance factors thus exposed the conflict that bootstrap support alone concealed.

Nuclear gene tree concordance at the node separating the Commersoniana group from *S. chacoense* (Tuberosa) confirms strong nuclear divergence, but the chloroplast tree recovers two *S. chacoense* clades that share plastid genes with the Commersoniana group. In the plastid tree, the *S. chacoense* North clade includes a strongly supported subclade comprising Spooner 6739 and Spooner & Salas 7204, while Ochoa 15584 was placed in more sister position the *S. chacoense* lineage. In contrast, the sister *S. chacoense* South clade is closely associated with the Commersoniana group in plastid tree, forming chloroplast-sharing clades.

ILS appeared at terminal branches in *S. commersonii* (Hawkes 706 × Okada 4583), *S. malmeanum* (Delhey s.n. × Okada 7303, and Brücher 18 × Brücher 16), and *S. chacoense* (Hawkes 3706 × Sleumer 3566), most likely from ancestral polymorphisms retained through recent, rapid divergence under weak reproductive isolation.

### Hybrid swarm structure in *S. commersonii* and *S. malmeanum*

Two accessions, *S. commersonii* (Boldrini & Boechat 264, **Fig. 8a**) and *S. malmeanum* (Delhey s.n., **Fig. 8b**), emerged as interspecific hybrids (*S. commersonii* × *S. malmeanum*), showing chloroplast-nuclear discordance and direct reticulation between the two Commersoniana lineages. *S. commersonii* (Boldrini & Boechat 264) likely originated as the result of a cross between *S. commersonii* (CST s.n.) and sister*S. malmeanum*, while *S. malmeanum* (Delhey s.n.) carries mixed ancestry fromsister *S. commersonii* and *S. malmeanum* lineages. *Solanum commersonii* (Okada 4583) shows both ILS and chloroplast capture.

**Fig. 8.**
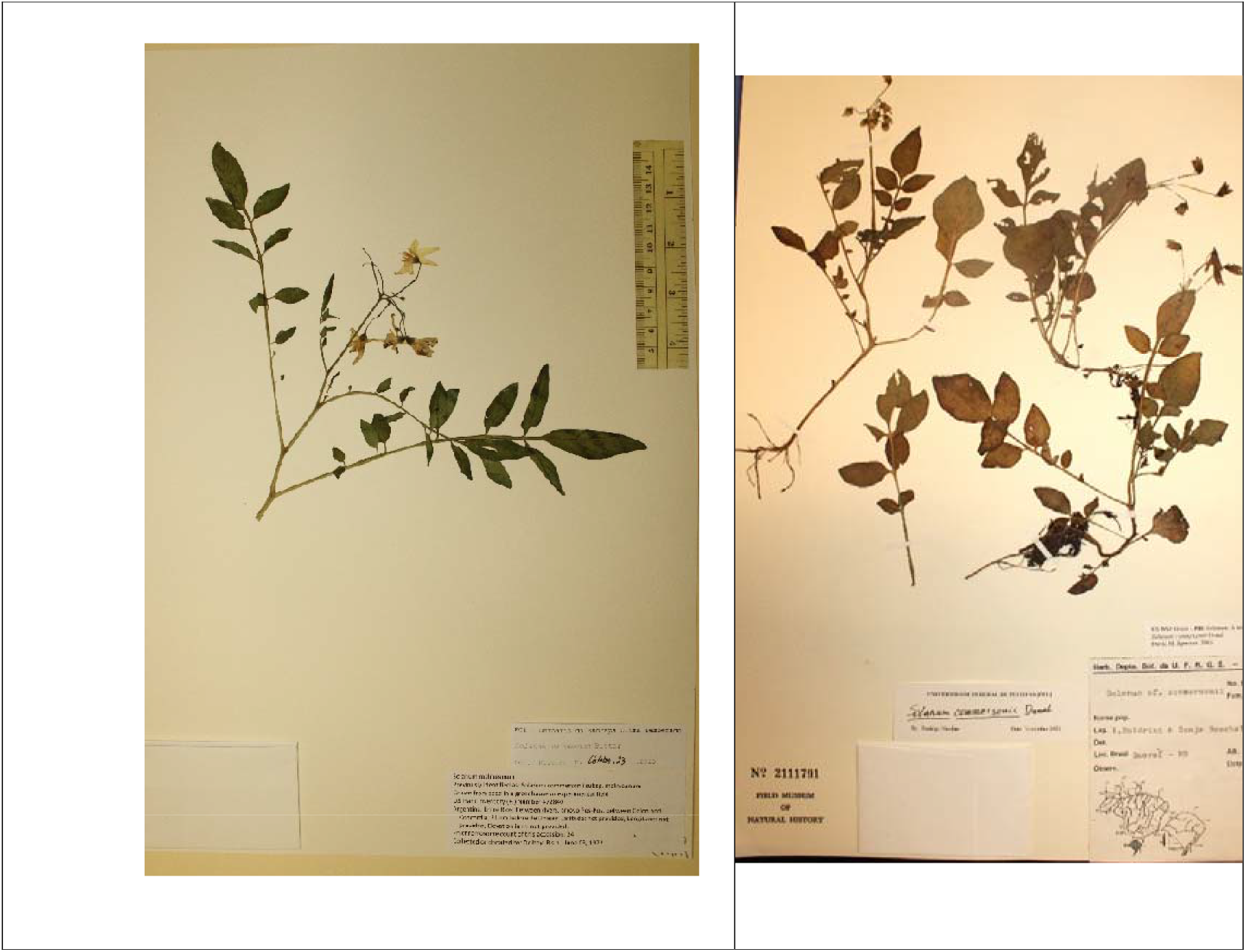
Hybrid swarm structure between *Solanum commersonii* and *S. malmeanum*. **a)** *S. commersonii* (Boldrini & Boechat 264); **b)** *S. malmeanum* (Delhey s.n.). Both accessions were recovered as interspecific hybrids (*S. commersonii* × *S. malmeanum*), exhibiting chloroplast–nuclear discordance and direct reticulation between the two Commersoniana lineages. *S. commersonii* (Boldrini & Boechat 264) likely originated from a cross between *S. commersonii* (CST s.n.) and a basal *S. malmeanum* lineage, whereas *S. malmeanum* (Delhey s.n.) shows mixed ancestry from basal lineages of both species.

### Deep reticulation between the Commersoniana group and *S. chacoense*

In *S. chacoense* North clade, accessions Spooner 6739 and Spooner & Salas 7204 reticulate with Ochoa 15584 and can be traced back to the *S. chacoense* node, signaling gene flow. In *S. chacoense* South, Correll 707A reticulates with sister nodes (Bordas s.n. and Okada 7309) and ancestral lineages (Sleumer 3566 and Hjerting 56). Deep reticulations linking the node at the base of the Commersoniana clade to *S. chacoense* point to ancient hybridization predating the reproductive barriers shaped by Endosperm Balance Number (EBN) and ecology **[11]**. Chloroplast capture from ancestral *S. chacoense* into the Commersoniana group, reinforced by repeated backcrossing, likely introgressed plastid genes that aided adaptation and diversification, and the simulation result (RF = 36.000, *p = 0.001*) rejects ILS as the sole process underlaying these deep conflicts.

#### Morphospace analysis

Vector fitting analysis (envfit) identified 16 morphological variables as significantly associated with species differentiation (p < 0.05). These were leaf length, leaf width, the leaf length-to-width ratio, length of the terminal leaflet lamina from its widest point to the apex, acumen length of the terminal leaflet, petiole length of the terminal leaflet, distance between lateral leaflets, length and width of the primary lateral leaflet lamina, width of the most distal lateral leaflet lamina, length from the widest point of the most distal lateral leaflet to the apex, length from the pedicel base to the articulation, and the shapes of the terminal leaflet, its apex, its base, and the base of the distal lateral leaflet.

Phylogenetic signal analysis indicated that 15 continuous morphological traits exhibited significant phylogenetic signal, including leaf width, leaf length-to-width ratio, length from the widest point of the terminal leaflet lamina to the apex, acumen length and petiole length of the terminal leaflet, distance between lateral leaflets, petiole length of the primary lateral leaflet, length and width of the most distal lateral leaflet lamina, length from the widest point of the most distal lateral leaflet to the apex, petiole length of the most distal lateral leaflet, number of lateral leaflets, calyx lobe length, length from the pedicel base to the articulation, and the ratio of the length from the pedicel base to the articulation to pedicel length. An NMDS on these variables (k = 2) reached a stress of 0.160, an acceptable two-dimensional fit to the morphological distances **[163, 164]**.

The ordination showed broad overlap among populations (**Fig. 9**). *S. commersonii*, *S. malmeanum*, and the putative hybrids (*S. commersonii* × *S. malmeanum*) were not clearly separated, and the southern *S. chacoense* (S) overlapped with them in part. Only the northern *S. chacoense* (N) formed a distinct, well-separated cluster.

**Fig. 9.**
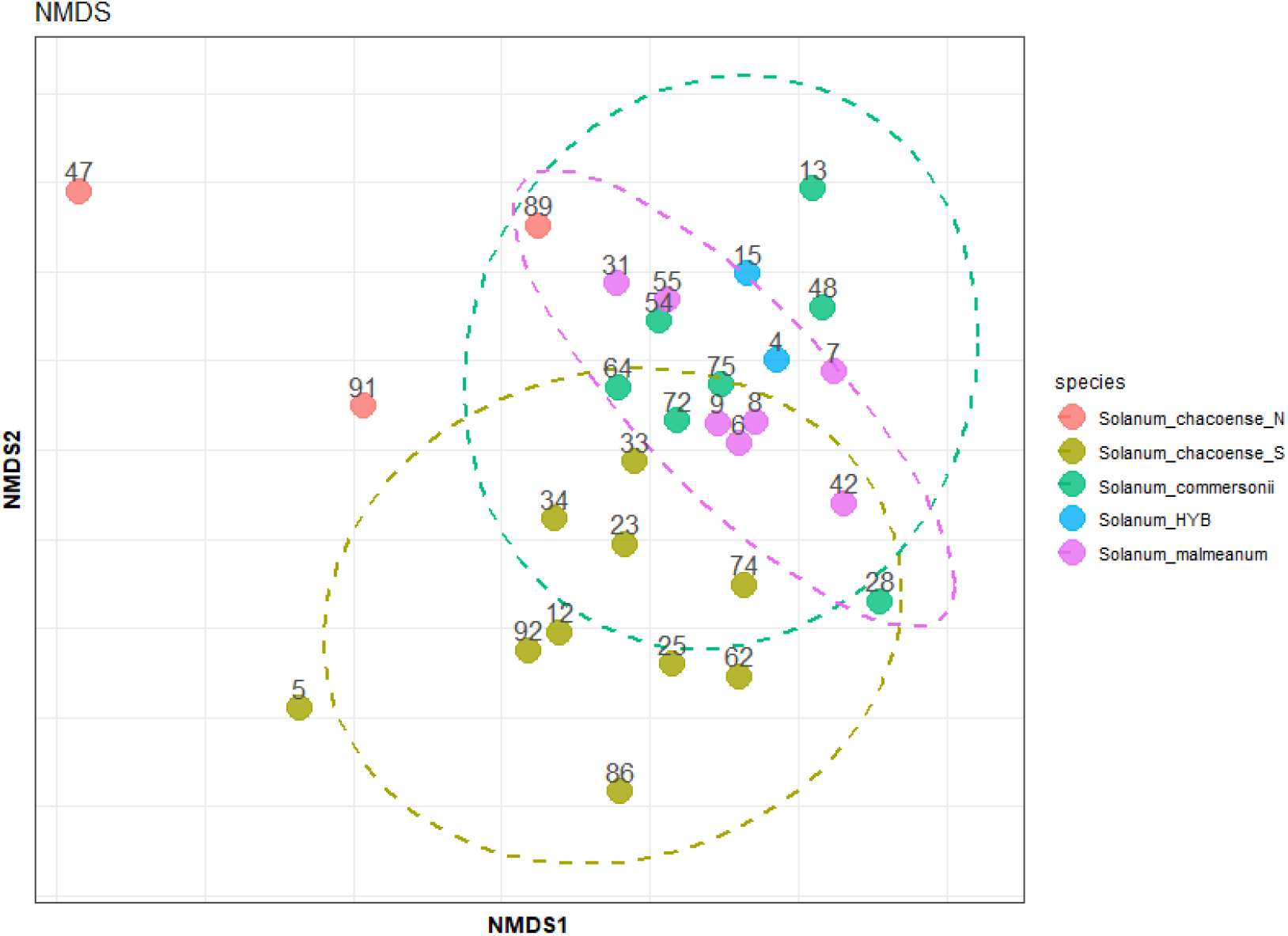
NMDS ordination of Southern South American wild potato populations from 16 significant morphological traits. Points are individual samples colored by species, *S. chacoense* N (red), *S. chacoense* S (olive green), *S. commersonii* (teal), the hybrids *S. commersonii* × *S. malmeanum* (blue), and *S. malmeanum* (lavender). Ellipses show 95% confidence intervals around group centroids.

#### Morphological variation of wild potato populations

We described four morphological groups, *S. chacoense* North, *S. chacoense* South, *S. commersonii*, *S. malmeanum*, and the interspecific hybrids *S. commersonii* (Boldrini & Boechat 264) and *S. malmeanum* (Delhey s.n.). Mean values and ranges (minimum– maximum) of continuous morphometric traits evaluated in Solanum chacoense (northern and southern populations), S. commersonii, S. malmeanum, and the hybrid (S. commersonii × S. malmeanum) are presented in in **Table S4** and discrete traits in **Table 2**.

**Table 2.**
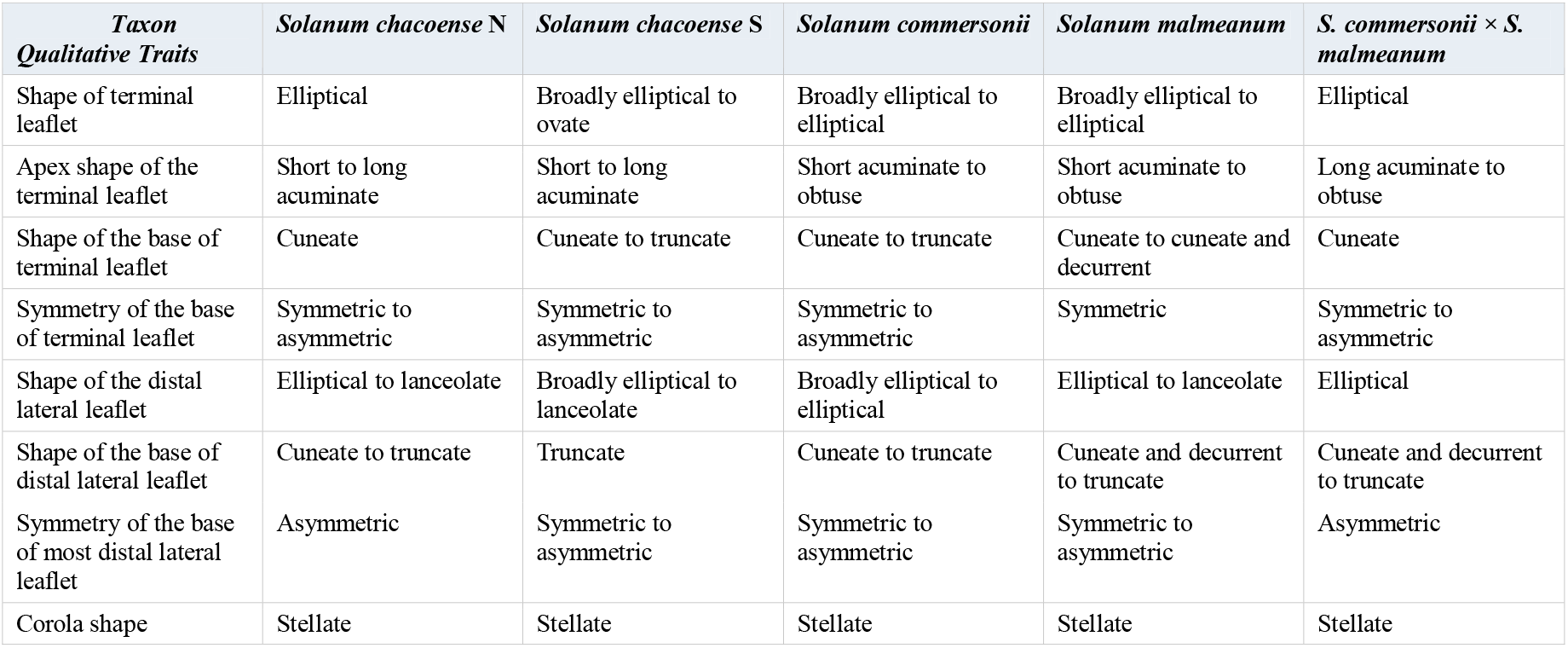
Discrete morphological traits of wild potato populations.

| Taxon | *Solanum chacoense* N | *Solanum chacoense* S | *Solanum commersonii* | *Solanum malmeanum* | *S. commersonii* × *S. malmeanum* |
| --- | --- | --- | --- | --- | --- |
| **Qualitative Traits** |  |  |  |  |  |
| Shape of terminal leaflet | Elliptical | Broadly elliptical to ovate | Broadly elliptical to elliptical | Broadly elliptical to elliptical | Elliptical |
| Apex shape of the terminal leaflet | Short to long acuminate | Short to long acuminate | Short acuminate to obtuse | Short acuminate to obtuse | Long acuminate to obtuse |
| Shape of the base of terminal leaflet | Cuneate | Cuneate to truncate | Cuneate to truncate | Cuneate to cuneate and decurrent | Cuneate |
| Symmetry of the base of terminal leaflet | Symmetric to asymmetric | Symmetric to asymmetric | Symmetric to asymmetric | Symmetric | Symmetric to asymmetric |
| Shape of the distal lateral leaflet | Elliptical to lanceolate | Broadly elliptical to lanceolate | Broadly elliptical to elliptical | Elliptical to lanceolate | Elliptical |
| Shape of the base of distal lateral leaflet | Cuneate to truncate | Truncate | Cuneate to truncate | Cuneate and decurrent to truncate | Cuneate and decurrent to truncate |
| Symmetry of the base of most distal lateral leaflet | Asymmetric | Symmetric to asymmetric | Symmetric to asymmetric | Symmetric to asymmetric | Asymmetric |
| Corolla shape | Stellate | Stellate | Stellate | Stellate | Stellate |

## Discussion

The A353 bait set resolves relationships at both family and genus levels, through its core loci **[165, 166]**, down to the more shallow, species levels, through the more variable flanking intronic regions captured alongside these core loci **[90, 91, 167, 168, 169, 170]**. Because they evolve faster and carry more variation from insertions and deletions (indels), supercontigs built from exons, introns, and flanking UTRs retain more variable sites than from alignments of exons alone **[171, 172]**, increasing their informativeness for resolving shallow relationships among recently diverged or rapidly diversifying lineages **[173, 174]**.

The nuclear topology recovered here differs from previous phylogenetic hypotheses, including Zhang et al. **[31]**, where *S. commersonii* was placed in Clade 4 South while *S. malmeanum* was recovered within Clade 3. Notably, the *S. malmeanum* accession PG4019 analyzed by Zhang et al. **[31]** was recovered within Clade 3 in their phylogeny. During our herbarium investigations, we verified that PG4019 is associated with the Okada 5074 collection, an accession included in our A353 target-capture dataset. However, whereas PG4019 was recovered within Clade 3, the Okada 5074 material analyzed here grouped with the remaining *S. malmeanum* accessions within the Commersoniana group. The conflicting placement of the same historical accession Okada 5074 following analysis in two distinct and independent phylogenomic datasets suggests that conflict may have resulted from sample processing, identification, or other biological artifacts. *Solanum malmeanum* is a southern South American potato species (Spooner et al. **[79]**; Nicolao et al. **[84]**, and its sister lineage (*S. commersonii*) has been placed in Clade 4 South in previous study by Yan et al. **[24]** . The placement of Okada 5074/PG4019 in Clade 3, a clade formed by northern South American potato species, is geographically and biologically incongruent. This suggests that the Clade 3 position is likely not representative of the genuine evolutionary placement of *S. malmeanum*, but rather reflects an artifact associated with the material analysed by Zhang et al. **[31].**

### Chloroplast capture rather than incomplete lineage sorting

Our PhyloFusion network, together with the coalescent simulations (observed RF distance = 36.000, *p = 0.001*), point to chloroplast capture rather than incomplete lineage sorting (ILS) as the main factor influencing cytonuclear discordance within the Commersoniana group (*S. commersonii* and *S. malmeanum*) and *S. chacoense* (Tuberosa group). Red edges (chloroplast capture) dominate the network **[Fig. 7]**, and the high measure of congruence between the chloroplast and nuclear species trees rejects ILS as the sole explanation for discordance **[59, 118, 163]**, pointing instead to introgressive hybridization that moved chloroplast genomes across species boundaries while nuclear genomes remained largely distinct. Disentangling such introgression from ILS is essential for interpreting cytonuclear discordance **[59]**, and our analyses were designed for exactly this distinction. Ancient hybridization, rather than non-sexual processes **[175]**, is the most plausible source of the deep chloroplast-nuclear conflicts we observe, although ILS still contributes at some terminal nodes.

Biparental inheritance of the nuclear genome versus the predominantly maternal inheritance of the plastid genome leads to a distinct phylogenetic signal from the two genomes, as well as different mutation rates and responses to drift and selection, complicating the reconstruction of species relationships **[114, 122, 192, 193, 194]**. Under some ecological conditions, this divergence can favour plastid capture outright, when an introgressed plastid haplotype confers an adaptive advantage such as improved photosynthetic performance or stress tolerance **[162, 195]**.

Hybridization is typically detected through gene tree discordance, the product of incongruent evolutionary histories among loci **[176, 177, 178]**. Chloroplast introgression through hybridization and backcrossing can fix the plastid genome of one lineage within another and pull its plastid phylogeny away from the nuclear species tree **[57, 161, 181],** whereas ILS arises in recently diverged species or large populations where ancestral polymorphisms persist long enough that rapid diversification cannot sort them completely **[179, 182, 183]**. Uniparental inheritance, limited recombination, and seed-mediated transmission further promote ILS **[161, 180]**. Such cytonuclear conflicts are well documented in plants, and reticulation generated by hybridization, introgression, ILS, and the interplay of sexual and asexual reproduction has long shaped the evolution of tuber-bearing *Solanum* **[12]**. Earlier work attributed *Solanum* incongruences to ILS and ancient hybridization **[184, 185]** and linked the high discordance in the genus to its rapid diversification **[186]**.

The interplay between hybridization and introgression (HI) and ILS reflects a temporal sequence. Gene flow mixed the genomes of two lineages before reproductive isolation was complete **[187]**, and the ILS that followed shows that, as the lineages diverged independently, some shared alleles, including introgressed ones, failed to sort cleanly during speciation **[57, 180]**. The result is a genome in which introgression introduces diversity, while ILS preserves variation shared between lineages that otherwise followed distinct evolutionary paths **[189, 190]**, a combination typical of recent adaptive radiations, where diversification into multiple ecological niches outpaces the establishment of reproductive barriers **[179, 183, 191]**.

### Nuclear signatures of chloroplast capture

The contrast between strong plastid discordance and weak nuclear D-statistics is itself informative. D-statistics interrogate only the nuclear genome, whereas the plastome generates the reticulations in our network and the cytonuclear discordance; thus, the absence of a strong nuclear footprint in the two hybrid accessions is consistent with chloroplast capture followed by repeated backcrossing, which restores the nuclear genome toward the recipient species while the captured plastome persists **[161, 162]**. *S. malmeanum* (Delhey s.n.) shows this most clearly. Analysis of the nuclear loci in Delhey s.n. support its identification as *S. malmeanum* and places it as sister to the Okada 5074 accession. In contrast, its plastome sequences show great similarity to *S. commersonii*. Thus, the phylogenetic discordance observed in Delhey s.n. is largely restricted to the plastid genome, with the nuclear genome retaining the expected *S. malmeanum* phylogenetic placement. *S. commersonii* (Boldrini & Boechat 264) is more ambiguous, because the nuclear species tree **[Fig. 3]** places it as sister to the entire *S. malmeanum* clade, hinting at historical gene flow, yet its D-statistic and per-window f*dM* are too weak to confirm recent introgression, again pointing to an older event partly erased by backcrossing.

The deepest event, capture from an ancestral *S. chacoense* lineage into the Commersoniana ancestor, lies beyond the reach of these tests by design. Because the introgression entered the common ancestor of *S. commersonii* and *S. malmeanum*, any resulting nuclear ancestry would be shared symmetrically by both species and therefore would be difficult to detect with a trio restricted to the Commersoniana group. The candidate donor accessions of southern *S. chacoense* (Bordas s.n. and Okada 7309) are nested tips within a single *S. chacoense* nuclear clade rather than a separate lineage that could be identified as a distinct donor taxon. The symmetric species-level result (D = 0.035, p = 0.63) is exactly what an ancient, shared chloroplast capture event would indicate. This deep-reticulation event is therefore supported by the plastome phylogeny, the PhyloFusion network, and the coalescent simulations rather than by the nuclear D-statistics, which neither confirm nor refute it. The nuclear genome of these wild potatoes appears largely sorted by species despite pervasive plastid sharing, the signature of chloroplast capture and backcrossing rather than recent, genome-wide hybridization.

### Morphology mirrors phylogenetic divergence

Morphology broadly supports these phylogenetic patterns. Strong phylogenetic signal in 18 continuous traits shows that leaf, petiole, and floral morphology are largely conserved, and pronounced differences in leaf size, leaflet dimensions, and acumen length between *S. chacoense* and the Commersoniana clade point to lineage-specific vegetative divergence. The NMDS ordination, despite overlap among *S. commersonii*, *S. malmeanum*, and the putative hybrids, separated the northern *S. chacoense* population as a distinct cluster. Leaf dimensions were intermediate in southern *S. chacoense* and smaller in *S. commersonii* and *S. malmeanum*. The leaf length to width ratio and petiole length follow the same conserved pattern.

The split of *S. chacoense* into northern and southern groups is supported by phylogenetic, biogeographic, and morphological evidence. Its nuclear and chloroplast phylogenies share a broadly similar topology despite both being polyphyletic, suggesting a shared history of geographic isolation and local adaptation. The moderate substructuring between northern and southern accessions likely reflects population differentiation, divergent plastomes, or ongoing speciation influenced by introgression and chloroplast capture. The nodes separating the two groups nonetheless carry low sCF and gCF, indicating weak phylogenetic support and substantial discordance attributable to chloroplast capture, as confirmed by the RF simulations, and the phylogenetic network places the two populations in distinct clusters with little reticulation between them.

### Hybridization and chloroplast capture within the Commersoniana group

Two chloroplast capture events link *S. chacoense* to the ancestor of the Commersoniana group, one from an ancestral sister node and the other from the southern *S. chacoense* populations (Bordas s.n. and Okada 7309). Repeated backcrossing after this introgression likely preserved the non-native plastid haplotypes while nuclear genomes continued to diverge **[161]**. Such introgression may also have contributed genetic variation that facilitated ecological diversification and niche colonization **[179].** This pattern is observed under adaptive radiation, where diversification proceeds alongside ongoing gene flow and incomplete reproductive isolation **[86, 191]**. This fits the broader picture of hybridization-driven speciation in *Solanum* **[179, 184, 185, 186]**, with reticulation supporting an origin of the Commersoniana group from ancestral *S. chacoense* through chloroplast capture and subsequent diversification, before pre- and post-zygotic barriers and ecological isolation were fully established.

*Solanum commersonii* and *S. malmeanum* (EBN = 1) are reproductively isolated from *S. chacoense* (EBN = 2) by their EBN difference **[86, 87]**, yet 2*n* gametes allow inter-EBN hybridization to occur **[72]**.

Reticulation between *S. malmeanum* (Delhey s.n.) and *S. commersonii* (Boldrini & Boechat 264) points to a hybrid swarm in which partially isolated lineages have exchanged genes, mainly through the plastome **[86, 87]**. Such introgressive gene flow can preserve genetic diversity and aid adaptation to heterogeneous environments, consistent with the rapid ecological diversification already documented in wild potatoes **[20]**. Morphologically, these hybrids are mostly intermediate between the parents, reflecting additive inheritance in leaf dimensions, petiole length, and floral traits.

However, some characters exhibit transgressive values: the ratio of terminal leaflet lamina length to width and the peduncle length exceed the range of both parents, whereas the distance from the widest point of the most distal lateral leaflet to the apex exceeds the maximum recorded for *S. commersonii*. These transgressive values point to genetic recombination and chloroplast capture as additional sources of morphological variation.

Pronounced reticulation among terminal branches in *S. chacoense* (Correll 707A, Hawkes 3706), *S. commersonii* (Okada 4583, Boldrini & Boechat 264), and *S. malmeanum* (Delhey s.n., Brücher 10, 16, 18, 19, and Okada 7309) points to ongoing gene flow within each group and reinforces the role of hybridization in their evolutionary dynamics **[79].** *Solanum commersonii* (Okada 4583) shows both ILS and chloroplast capture. Its reticulation with Hawkes 706 reflects ILS from unsorted ancestral polymorphism, whereas its plastid affinity with Okada 5059 marks chloroplast capture through hybridization and backcrossing. Reticulations among six *S. malmeanum* accessions (Delhey s.n., Brücher 10, 16, 18, and 19, and Okada 7309) (Figure 1) describe a hybrid swarm. Okada 7303 and Brücher 18 carry both red (chloroplast capture) and blue (ILS) edges, a mixed zone where ancestral nuclear polymorphism persists alongside introgressed plastids. The other four show chloroplast capture only, consistent with older hybridization, after which nuclear sorting erased detectable ILS. Recurrent backcrossing or selection favoring the captured haplotypes could cause this, particularly where plastid genes aid thermotolerance or photosynthesis. Together, these clustered reticulations support a hybrid swarm in *S. malmeanum*, with partially isolated lineages exchanging both nuclear and chloroplast material.

Distinct environmental mechanisms can promote reproductive isolation, stabilizing hybrid lineages in divergent habitats while limiting hybridization where habitats do not diverge **[198]**, and the differing inheritance and drift of plastid versus nuclear genomes further complicate these dynamics while enhancing adaptation to diverse niches, as seen in other angiosperms **[51, 79, 122, 192]**.

### Breeding implications

For pre-breeding, the strong phylogenetic signal in leaf, petiole, and floral traits means that morphological characters are evolutionarily conserved and predictable, and these can be reliably selected for when choosing parental lines. The *S. commersonii* × *S. malmeanum* hybrids were mostly intermediate, reflecting additive inheritance, but the transgressive values seen in the leaflet length to width ratio, peduncle length, and distal leaflet apex length represent phenotypic variation beyond either parent’s range, variation that breeding programs could exploit for new combinations of leaf architecture, floral structure, or adaptive traits in cultivated germplasm.

Variation in reproductive traits, such as peduncle, pedicel, anther, and style length, points to genotypes with favorable flowering and cross-compatibility for introgressing wild alleles into diploid or tetraploid cultivated backgrounds. Tuber traits are equally important agronomically but are absent from herbarium material, and thus, extending this approach to germplasm accessions would allow tuber size, shape, yield, and disease resistance to be assessed alongside the morphological and reproductive traits considered here.

### Species boundaries and speciation within Commersoniana

Speciation within Commersoniana likely involved ancient geographic isolation together with episodic drift and gene flow. The taxonomic status of *S. malmeanum* remains debated **[2, 3, 6, 20]** since it closely resembles *S. commersonii* in morphology, with stellate flowers and small tubers, yet differs ecologically in ways that point to local adaptation **[20]**, and AFLP data have supported genetic differentiation between *S. commersonii* subsp. *commersonii* and *S. commersonii* subsp. *malmeanum* **[199]**.

Within Commersoniana, *S. malmeanum* has been proposed as a homoploid hybrid of *S. commersonii* (EBN = 1) and *S. chacoense* (EBN = 2) that retained the diploid state (2n = 2x = 24) **[2, 86]**. This hypothesis implies gene flow across overlapping ranges, mediated by such pollinators as *Bombus* spp. capable of flying up to 1750 m **[192, 193]** and facilitated by 2*n* gametes **[72, 86, 88]**. However, the symmetric, non-significant species-level D-statistic gives no sign of the asymmetric *S. chacoense* contribution that this scenario predicts; thus, either *S. malmeanum* is not of recent homoploid hybrid origin or the nuclear signal has since been erased by backcrossing and lineage sorting.

Species differentiation here is not caused by geographic isolation alone but by the interplay of conserved versus plastic morphology and a genome that remains permeable to hybridization. The departure of some morphological characters from classical models of evolution reflects the pervasive influence of introgression throughout the history of this group.

## Conclusions

Our results show that recurrent hybridization and cytonuclear discordance shape the evolutionary history of Southern South American wild potatoes. The Commersoniana group may have originated through reticulation with an ancestral *S. chacoense* population, as evidenced by the chloroplast capture detected between them, consistent with a sympatric mode of speciation in which a reproductive barrier developed within that ancestral *S. chacoense* population while still in contact with its descendants. The relationship between *S. commersonii* and *S. malmeanum* instead fits a parapatric model, in which a Commersoniana putative ancestral population expanded into adjacent but ecologically distinct niches giving rise to the two species. Although both species share an EBN of 1, indicating potential post-zygotic compatibility, their clear nuclear differentiation suggests that they represent distinct evolutionary lineages despite the potential for interspecific gene flow. *Solanum commersonii* tends to occur in the Eastern side of the Commersoniana area of distribution, while *S. malmeanum* is more common along the Western side, existing an overlapping area of distribution, that may have originated or increased in area more recently. Across all analyses, the plastome carries the strongest signal of this reticulate history, whereas the nuclear genome remains largely sorted by species, indicating that chloroplast capture followed by backcrossing, rather than recent genome-wide hybridization, has shaped the group.

Several questions remain beyond the reach of our current data. Whole-genome or denser SNP datasets, rather than the target-capture loci used here, would provide the power to confirm the weak nuclear signals in single accessions such as *S. commersonii* (Boldrini & Boechat 264). Further, methods that recover the direction and timing of gene flow, such as DFOIL, HyDe, or multispecies-coalescent-with-introgression models in PhyloNet and BPP, would test whether the deep *S. chacoense* to Commersoniana capture reflects a single ancestral event or several independent ones. Sequencing the mitochondrial genome alongside the plastome would show whether both cytoplasmic compartments were co-captured, strengthening the inference of maternal capture over chance plastid sharing, while broader and more dense population sampling, especially of northern and southern *S. chacoense* and the rarer *S. commersonii* accessions, would enable allele-frequency methods, such as f-branch and population-level Patterson’s D, to localize introgression on specific branches. Nonetheless, our study provides a robust application of Angiosperms353 to reconstructing phylogenetic relationships at shallow scales and leveraging these loci to detect and trace the putative source of reticulation events. The results also demonstrate the potential use of Angiosperms353 for the study of plant genetic resources. Finally, dated phylogenetic and demographic modelling would place these capture and backcrossing events in time.

## Acknowledgments

R.N. acknowledges Universidade Federal de Pelotas, Embrapa Clima Temperado, the Botanical Research Institute of Texas (BRIT), the University of Helsinki/LUOMUS, and CAPES (88887.622551/2021-00 and PDSE 88881.846409/2023-01) for supporting his PhD research. R.N. also acknowledges the International Association for Plant Taxonomy (IAPT) 2022, Bill Dahl Graduate Student Research Award, the Botanical Society of America (BSA) 2023 award, and the LinnéSys: Systematics Research Fund from the Linnean Society of London and the Systematics Association (2024) for research grants that supported herbarium reviews and DNA sequencing. R.N. acknowledges the Herbarium of Wisconsin University, Field Museum, and BRIT for leaf samples. GH acknowledges Conselho Nacional de Desenvolvimento Científico e Tecnológico – CNPq for the research productivity grant (312897/2025-1). We acknowledge the open-access support of Helsinki University Library. The CSC – IT Center for Science, Finland, provided free access to high-performance computing resources essential for conducting this research. We are grateful for support from the TFK – MOMENT project and the EDUFI Fellowship program.

This research represents a partial fulfilment of the requirements for the degree of Doctor of Philosophy (PhD) at the Universidade Federal de Pelotas (UFPel) in collaboration with Embrapa Clima Temperado, Brazil by R.N.

We also thank Carol Ann Pelli for editing the manuscript.

## Authors’ contributions

**Conceptualization:** R.N., P.P., M.G., G.H. **Funding acquisition**: R.N., P.P., M.G., C.M.C., G.H. **Supervision**: P.P., M.G., C.M.C., G.H. **Formal analysis**: R.N., P.P., R.D. **Investigation**: R.N., P.P. **Methodology**: R.N., P.P., R.D. **Project administration**: R.N., G.H., C.M.C. **Resources**: R.N., P.P., M.B., M.G., G.H. **Software**: R.N., P.P., R.D. **Validation**: P.P., R.D., M.B., M.G., C.M.C. G.H. **Visualization**: R.N., P.P., G.H. **Writing – original draft:** R.N. **Writing – review & editing:** P.P., M.G., C.M.C., G.H.

## Declarations

### Data availability

All data are available within the manuscript and its supplementary files, or through the Open Science Framework (OSF) at https://osf.io/mdsqj.

## Footnotes

1 The EBN explains why some crosses between potato species succeed and others fail even when the parents share the same chromosome number or ploidy. Successful seed set depends not on ploidy alone but on a genetic balance between maternal and paternal contributions to the endosperm. Each species carries an EBN, a numerical value reflecting the effective ploidy of its genome for endosperm compatibility. The crucial requirement is a 2 to 1 maternal-to-paternal EBN ratio in the endosperm. When that ratio holds, the cross usually works; when it is off, the endosperm fails and the seed aborts **[13]**. Reported ploidy and EBN combinations in potatoes are 6x (4EBN), 4x (4EBN), 4x (2EBN), 2x (2EBN), and 2x (1EBN). The EBN theory **[13]** capped several decades of cytogenetic and embryological work on *Solanum* and other genera. Von Wangenheim **[14]** linked chromosome number to crossability (viability of offspring resulting from crosses), showing that ploidy differences often raise hybridization barriers expressed mainly as seed failure, while von Wangenheim, Peloquin, and Hougas **[15]** traced haploid formation in *Solanum tuberosum* and found that abnormal endosperm development reliably preceded embryo abortion in incompatible crosses. In wheat, Gill and Waines **[16]** showed that genomic imprinting, rather than chromosome number alone, can govern endosperm success in interspecific crosses of *Triticum* spp. L. and *Aegilops* spp. L., where compatibility hinges on the correct maternal-to-paternal genomic ratio.

2 The study of potato diversity and its role in global agriculture owes much to the foundational work of John Gregory (Jack) Hawkes (1915-2007), Donovan S. Correll (1908-1983), Heintz Brücher (1915-1991), Carlos M. Ochoa (1920-2008), David Michael Spooner (1949-2022), Jens Peter Knudsen Hjerting (1917-2012), and Katsuo Armando Okada (1935-2014). Their contributions in taxonomy, systematics, and genetic resources conservation bridged traditional botanical exploration with emerging paradigms in genetic resource management, breeding, and evolutionary biology. Hawkes, a pioneer of in situ and ex situ germplasm conservation, led extensive Andean collecting expeditions and assembled one of the most comprehensive *Solanum* germplasm collections, and his monographs synthesized morphology, geography, and cytogenetics into the first coherent overview of wild and cultivated potato diversity. Brücher advanced our understanding of cytology, reproductive biology, and hybridization under challenging circumstances, linking these insights to breeding potential. Ochoa combined exceptional field expertise with rigorous taxonomy, describing numerous species and documenting indigenous cultivation systems. Hjerting linked wild species research to cultivar development through cytogenetics and breeding, while Okada elucidated chromosome behaviour and genetic compatibility in *Solanum* sect. *Petota*. Spooner introduced molecular systematics to potato research, integrating DNA-based phylogenetics with morphology and ecology to produce a modern, evolutionarily informed classification. Their combined legacies include herbarium collections, germplasm bank accessions, and reference works that underpin potato taxonomy and global food security. These resources have proven crucial for identifying resistance genes to late blight, viruses, and abiotic stresses, assets that grow more important as climate change, pathogens, and environmental instability threaten production. Their work continues to inform agriculture, ecology, and evolution, and by documenting agrobiodiversity they provided baselines to measure genetic erosion, tools to breed climate-adapted cultivars, and insights into domestication. Their legacy shows how meticulous natural history, embedded in global conservation networks, can become a cornerstone of sustainable agriculture, ecological resilience, and economic stability.

