## Supplementary Material for "Reticulate History of Southern South American Wild Potatoes: Recurrent Hybridization, Cytonuclear Discordance, and Phylogenetic Placement of *Solanum malmeanum*"

**Geographical distribution of potato sampling**

**Table S1.** Geographic, taxonomic, and herbarium metadata for sampled wild potato *Solanum* accessions. Information provided for each accession includes species code (sample number), taxonomic identity based on morphological circumscription for herbarium-derived specimens, collector and voucher number, herbarium acronym or source study, geographic coordinates (latitude and longitude; UTM), and collection date.

| **Code** | **Potato species** | **Collector / number** | **Latitude** | **Longitude** | **Herbarium / study** |
| --- | --- | --- | --- | --- | --- |
| 1 | *Solanum chacoense* Bitter | Ochoa 15584 | -18.67 | -65.17 | F |
| 2 | *Solanum chacoense* Bitter | Spooner & Salas 7204 | -14.22 | -69.45 | WIS |
| 3 | *Solanum chacoense* Bitter | Spooner 6739 | -16.4 | -67.94 | WIS |
| 4 | *Solanum chacoense* Bitter | Bordas S/N | -23.1 | -55.92 | WIS |
| 5 | *Solanum chacoense* Bitter | Correll 707A | -27.25 | -66.25 | WIS |
| 6 | *Solanum chacoense* Bitter | Hawkes 3196 | -32.57 | -65 | WIS |
| 7 | *Solanum chacoense* Bitter | Hawkes 3706 | -25.17 | -65.83 | WIS |
| 8 | *Solanum chacoense* Bitter | Hjerting 56 | -27.25 | -65.87 | WIS |
| 9 | *Solanum chacoense* Bitter | Hjerting 6377 | -27.43 | -66 | WIS |
| 10 | *Solanum chacoense* Bitter | Okada 5628 | -25.15 | -65.73 | WIS |
| 11 | *Solanum chacoense* Bitter | Sleumer 3566 | -23.92 | -65.42 | WIS |
| 12 | *Solanum commersonii* Poir. | Hawkes 706 | -37.75 | -58.3 | WIS |
| 13 | *Solanum commersonii* Poir. | Okada 4583 | -37.73 | -58.3 | WIS |
| 14 | *Solanum commersonii* Poir. | Okada 5059 | -37.87 | -58.25 | WIS |
| 15 | *Solanum commersonii* Poir. | Okada 7309 | -27.55 | -58.53 | WIS |
| 16 | *Solanum commersonii* Poir. | Boldrini & Boechát 264 | -30.38 | -56.45 | F |
| 17 | *Solanum commersonii* Poir. | CST S/N | -31.77 | -52.33 | WIS |
| 18 | *Solanum commersonii* Poir. | FB 4001 | -32.2206 | -57.4625 | Zhang et al. 2025 |
| 19 | *Solanum malmeanum* Bitter | Delhey S/N | -32.22 | -58.33 | WIS |
| 20 | *Solanum malmeanum* Bitter | Brücher 10 | -29.9 | -60.3 | WIS |
| 21 | *Solanum malmeanum* Bitter | Brücher 16 | -28.38 | -53.92 | WIS |
| 22 | *Solanum malmeanum* Bitter | Brücher 18 | -23 | -60 | WIS |
| 23 | *Solanum malmeanum* Bitter | Brücher 19 | -28.38 | -53.92 | WIS |
| 24 | *Solanum malmeanum* Bitter | Hawkes 725 | -31.39 | -57.32 | WIS |
| 25 | *Solanum malmeanum* Bitter | Krapovickas & Cristóbal 14792 | -28.36 | -56.13 | WIS |
| 26 | *Solanum malmeanum* Bitter | Okada 5074 | -29.72 | -57.58 | WIS |
| 27 | *Solanum malmeanum* Bitter | Okada 7254 | -33.03 | -58.53 | WIS |
| 28 | *Solanum malmeanum* Bitter | Okada 7303 | -27.32 | -58.9 | WIS |
| 29 | *Solanum malmeanum* Bitter | Okada 7313 | -31.77 | -60.53 | WIS |
| 30 | *Solanum malmeanum* Bitter | Okada 5074 | -29.5667 | -57.5333 | Zhang et al. 2025 |
| 31 | *Solanum immite* Dunal | Ochoa 13346 | -8.1167 | -79.0333 | Zhang et al. 2025 |
| 32 | *Solanum albornozii* Correll | Castillo et al. 5032 | -4 | -79.2669 | Zhang et al. 2025 |
| 33 | *Solanum acroglossum* Juz. | Ochoa 11297 | -10.4672 | -76.3478 | Zhang et al. 2025 |
| 34 | *Solanum mochiquense* Ochoa | GLKS 127/1 | -11.35 | -77.3833 | Zhang et al. 2025 |
| 35 | *Solanum acroscopicum* Ochoa | Ochoa 81 | -15.2481 | -72.86 | Zhang et al. 2025 |
| 36 | *Solanum cajamarquense* Ochoa | Ochoa 16118 | -7.3356 | -78.1703 | Zhang et al. 2025 |
| 37 | *Solanum infundibuliforme* Phil. | Fernandez et al. 6539 | -19.8333 | -65.7 | Zhang et al. 2025 |
| 38 | *Solanum infundibuliforme* Phil. | Ochoa 11941 | -19.6333 | -65.2167 | Zhang et al. 2025 |
| 39 | *Solanum piurae* Bitter | Ochoa 13959 | -5.3119 | -79.6258 | Zhang et al. 2025 |
| 40 | *Solanum huancabambense* Ochoa | Ochoa 11627 | -5.2942 | -79.4836 | Zhang et al. 2025 |
| 41 | *Solanum verrucosum* Schltdl. | Hjerting 7394 | 24.87 | -100.26 | Zhang et al. 2025 |
| 42 | *Solanum okadae* Hawkes & Hjerting | Okada 7657 | -25.1833 | -65.85 | Zhang et al. 2025 |
| 43 | *Solanum microdontum* Bitter | Hoopes & Okada 23 | -17.6667 | -65.3 | Zhang et al. 2025 |
| 44 | *Solanum kurtzianum* Bitter & Wittm. | Okada 4986 | -29.4167 | -68 | Zhang et al. 2025 |
| 45 | *Solanum flahaultii* Bitter | Lopez S/N | 5.13333 | -74.05 | Zhang et al. 2025 |
| 46 | *Solanum paucissectum* Ochoa | Ochoa 14818 | -5.3981 | -79.4597 | Zhang et al. 2025 |

**Table S2** Vegetative and reproductive characters evaluated.

| **No.** | **Trait** | **Organ** | **Type (Continuous/Discrete)** |
| --- | --- | --- | --- |
| 1 | Leaf length | Leaf | Continuous |
| 2 | Leaf width | Leaf | Continuous |
| 3 | Ratio: leaf length/width | Leaf | Continuous |
| 4 | Length from widest point of leaf to apex | Leaf | Continuous |
| 5 | Length of terminal leaflet lamina | Terminal leaflet | Continuous |
| 6 | Width of terminal leaflet lamina | Terminal leaflet | Continuous |
| 7 | Ratio: terminal leaflet lamina length/width | Terminal leaflet | Continuous |
| 8 | Length from widest point of terminal leaflet lamina to apex | Terminal leaflet | Continuous |
| 9 | Acumen length of terminal leaflet lamina | Terminal leaflet | Continuous |
| 10 | Petiole length of terminal leaflet | Terminal leaflet | Continuous |
| 11 | Distance between lateral leaflets | Leaf | Continuous |
| 12 | Length of primary lateral leaflet lamina | Primary lateral leaflet | Continuous |
| 13 | Width of primary lateral leaflet lamina | Primary lateral leaflet | Continuous |
| 14 | Ratio: primary lateral leaflet lamina length/width | Primary lateral leaflet | Continuous |
| 15 | Length from widest point of primary lateral leaflet lamina to apex | Primary lateral leaflet | Continuous |
| 16 | Petiole length of primary lateral leaflet | Primary lateral leaflet | Continuous |
| 17 | Length of most distal lateral leaflet lamina | Most distal lateral leaflet | Continuous |
| 18 | Width of most distal lateral leaflet lamina | Most distal lateral leaflet | Continuous |
| 19 | Ratio: most distal lateral leaflet lamina length/width | Most distal lateral leaflet | Continuous |
| 20 | Length from widest point of most distal lateral leaflet lamina to apex | Most distal lateral leaflet | Continuous |
| 21 | Petiole length of most distal lateral leaflet | Most distal lateral leaflet | Continuous |
| 22 | Number of lateral leaflets | Leaf | Discrete |
| 23 | Number of interstitial leaflet segments between lateral leaflets | Leaf | Discrete |
| 24 | Shape of terminal leaflet | Terminal leaflet | Discrete |
| 25 | Shape of apex of terminal leaflet | Terminal leaflet | Discrete |
| 26 | Shape of base of terminal leaflet | Terminal leaflet | Discrete |
| 27 | Symmetry of base of terminal leaflet | Terminal leaflet | Discrete |
| 28 | Shape of distal lateral leaflet | Distal lateral leaflet | Discrete |
| 29 | Shape of base of distal lateral leaflet | Distal lateral leaflet | Discrete |
| 30 | Symmetry of base of most distal lateral leaflet | Most distal lateral leaflet | Discrete |
| 31 | Peduncle length | Inflorescence | Continuous |
| 32 | Pedicel length | Pedicel | Continuous |
| 33 | Length from base of pedicel to articulation | Pedicel | Continuous |
| 34 | Ratio: length from base of pedicel to articulation/pedicel length | Pedicel | Continuous |
| 35 | Calyx lacinia length | Calyx | Continuous |
| 36 | Length of calyx acumen | Calyx | Continuous |
| 37 | Total calyx length | Calyx | Continuous |
| 38 | Calyx lobe length | Calyx lobe | Continuous |
| 39 | Calyx lobe width | Calyx lobe | Continuous |
| 40 | Length of sepal lobe | Sepal | Continuous |
| 41 | Radius of corolla lobe | Corolla lobe | Continuous |
| 42 | Length of corolla lobe from base to apex | Corolla lobe | Continuous |
| 43 | Width of corolla lobe at base | Corolla lobe | Continuous |
| 44 | Acumen length of corolla lobe | Corolla lobe | Continuous |
| 45 | Anther length | Anther | Continuous |
| 46 | Style length | Style | Continuous |
| 47 | Style length exceeding stamens | Style | Continuous |
| 48 | Corolla shape | Corolla | Discrete |

**Supplementary Method 1**

**Phylogenetic signal of continuous traits**

We mapped 38 continuous traits onto the MCC tree using the ‘ContMap’ function in the {phytools} package^[1]^ in RStudio version 2025.5.0^[2]^. Ancestral character states were estimated using an ML-based procedure under the assumption that characters evolve under a Brownian motion model. To evaluate whether the observed trait similarities reflect shared evolutionary history, we assessed the phylogenetic signal of continuous characters using Blomberg’s *K* statistic^[3],^^[4]^, calculated using the ‘phylosig’ function of the {phytools} package^5^ in RStudio version 2025.5.0^[5]^.

**Gene recovery with HybPiper**

***Nuclear genes (Angiosperms353)***

Read counts ranged from 81 059 in *Solanum chacoense* (Hawkes 3706) to 1 083 016 in *Solanum malmeanum* (Delhey s.n.), and four samples exceeded one million reads (**Table S1**). Mapped Angiosperms353 reads spanned 27 160 in *S. malmeanum* (Brücher 10) to 532 188 in *S. chacoense* (Correll 707A). Genes recovered ranged from 246 in *S. stoloniferum* (Waterfall & Wallis 13489) to 347 in *S. chacoense* (Correll 707A), and genes at 75% coverage or more from 36 in that same *S. stoloniferum* accession to 284 in *S. malmeanum* (Okada 5074). We identified and removed 26 paralogous nuclear genes.

**Table S3.** Recovery statistics from nuclear target-capture sequencing using the Angiosperms353 probe set.

| **Species** | **Collector** | **Herbarium** | **Number of reads** | **Reads mapped** | **% on Target** | **Genes with sequences** | **Genes at 75%** |
| --- | --- | --- | --- | --- | --- | --- | --- |
| *Solanum chacoense* Bitter | Bordas E., s.n. | WIS | 474 675 | 251 378 | 53 | 343 | 230 |
| *Solanum chacoense* Bitter | Correll D.S., 707A | WIS | 1 012 931 | 532 188 | 52.5 | 347 | 283 |
| *Solanum chacoense* Bitter | Hawkes J.G. et al., 3196 | WIS | 372 020 | 193 435 | 52 | 341 | 228 |
| *Solanum chacoense* Bitter | Hawkes J.G. et al., 3706 | WIS | 81 059 | 37 365 | 46.1 | 254 | 40 |
| *Solanum chacoense* Bitter | Hjerting J.P., 56 | WIS | 236 096 | 86 703 | 36.7 | 327 | 156 |
| *Solanum chacoense* Bitter | Hjerting J.P., 6377 | WIS | 623 661 | 272 067 | 43.6 | 345 | 272 |
| *Solanum chacoense* Bitter | Ochoa C., 15584 | F | 836 285 | 311 852 | 37.3 | 345 | 276 |
| *Solanum chacoense* Bitter | Okada K.A., 5628 | WIS | 314 609 | 79 310 | 25.2 | 329 | 178 |
| *Solanum chacoense* Bitter | Okada K.A., 7309 | WIS | 131 277 | 61 842 | 47.1 | 300 | 77 |
| *Solanum chacoense* Bitter | Sleumer H., 3566 | WIS | 848 436 | 370 183 | 43.6 | 347 | 271 |
| *Solanum chacoense* Bitter | Spooner D.M. & Salas A., 7204 | WIS | 460 312 | 203 780 | 44.3 | 342 | 219 |
| *Solanum chacoense* Bitter | Spooner D.M. et al., 6379 | WIS | 276 892 | 116 625 | 42.1 | 336 | 226 |
| *Solanum chacoense* Bitter | Tressens 4742 | F | 379 803 | 158 678 | 41.8 | 332 | 179 |
| *Solanum commersonii Poir.* | CST, s.n. | WIS | 1 057 490 | 467 019 | 44.2 | 345 | 274 |
| *Solanum commersonii Poir.* | Boldrini I. & Boechat S., 264 | F | 246 872 | 62 795 | 25.4 | 318 | 183 |
| *Solanum commersonii Poir.* | Hawkes J.G. et al., 706 | WIS | 162 288 | 85 744 | 52.8 | 324 | 114 |
| *Solanum commersonii Poir.* | Okada K.A., 4583 | WIS | 438 364 | 231 021 | 52.7 | 341 | 226 |
| *Solanum commersonii Poir.* | Okada K.A., 5059 | WIS | 406 536 | 192 401 | 47.3 | 343 | 229 |
| *Solanum malmeanum* Bitter | Brücher E.H., 10 | WIS | 832 765 | 27 160 | 3.3 | 276 | 141 |
| *Solanum malmeanum* Bitter | Brücher E.H., 16 | WIS | 702 991 | 347 371 | 49.4 | 345 | 265 |
| *Solanum malmeanum* Bitter | Brücher E.H., 18 | WIS | 312 294 | 77 243 | 24.7 | 330 | 182 |
| *Solanum malmeanum* Bitter | Brücher E.H., 19 | WIS | 307 622 | 130 959 | 42.6 | 332 | 195 |
| *Solanum malmeanum* Bitter | Delhey R., s.n. | WIS | 1 083 016 | 398 432 | 36.8 | 346 | 268 |
| *Solanum malmeanum* Bitter | Hawkes J.G. et al., 725 | WIS | 189 964 | 70 607 | 37.2 | 311 | 100 |
| *Solanum malmeanum* Bitter | Krapovickas A. & Cristóbal C.L., 14792 | WIS | 322 990 | 155 388 | 48.1 | 334 | 143 |
| *Solanum malmeanum* Bitter | Okada K.A., 5074 | WIS | 1 027 545 | 450 284 | 43.8 | 346 | 284 |
| *Solanum malmeanum* Bitter | Okada K.A., 7254 | WIS | 591 762 | 276 948 | 46.8 | 344 | 260 |
| *Solanum malmeanum* Bitter | Okada K.A., 7303 | WIS | 523 353 | 217 509 | 41.6 | 344 | 257 |
| *Solanum malmeanum* Bitter | Okada K.A., 7313 | WIS | 137 904 | 37 742 | 27.4 | 275 | 71 |
| *Solanum stoloniferum* Schltdl. | Waterfall U.T. & Wallis C.S., 13489 | BRIT | 164 878 | 32 660 | 19.8 | 246 | 36 |

Values are given per accession for *S. chacoense*, *S. commersonii*, *S. malmeanum*, and *S. stoloniferum*, with collector and herbarium, total reads, reads mapped to the reference, percentage on target, Angiosperms353 genes recovered, and genes recovered at 75% or more of target length.

***Chloroplast genes***

Recovered chloroplast genes ranged from 5 (*S. chacoense*, Hawkes 3706 and Spooner 6739) to 74 (*S. malmeanum*, Delhey s.n.) (Table S2).

**Table S2.** Recovery statistics of chloroplast sequences from off-target capture with the Angiosperms353 probe set.

| **Species** | **Collector** | **Herbarium** | **Number of reads** | **Reads mapped** | **% on Target** | **Genes with sequences** | **Genes at 75%** |
| --- | --- | --- | --- | --- | --- | --- | --- |
| *Solanum chacoense* Bitter | Bordas E., s.n. | WIS | 474 675 | 8 362 | 1.8 | 58 | 54 |
| *Solanum chacoense* Bitter | Correll D.S., 707A | WIS | 1 012 931 | 4 047 | 0.4 | 46 | 39 |
| *Solanum chacoense* Bitter | Hawkes J.G. et al., 3196 | WIS | 372 020 | 9 940 | 2.7 | 65 | 63 |
| *Solanum chacoense* Bitter | Hawkes J.G. et al., 3706 | WIS | 81 059 | 571 | 0.7 | 5 | 0 |
| *Solanum chacoense* Bitter | Hjerting J.P., 56 | WIS | 236 096 | 5 328 | 2.3 | 59 | 53 |
| *Solanum chacoense* Bitter | Hjerting J.P., 6377 | WIS | 623 661 | 7 499 | 1.2 | 61 | 59 |
| *Solanum chacoense* Bitter | Ochoa C., 15584 | F | 836 285 | 5 452 | 0.7 | 51 | 47 |
| *Solanum chacoense* Bitter | Okada K.A., 5628 | WIS | 314 609 | 8 242 | 2.6 | 64 | 60 |
| *Solanum chacoense* Bitter | Okada K.A., 7309 | WIS | 131 277 | 3 278 | 2.5 | 39 | 28 |
| *Solanum chacoense* Bitter | Sleumer H., 3566 | WIS | 848 436 | 8 024 | 0.9 | 67 | 61 |
| *Solanum chacoense* Bitter | Spooner D.M. & Salas A., 7204 | WIS | 46 0312 | 3 345 | 0.7 | 31 | 19 |
| *Solanum chacoense* Bitter | Spooner D.M. et al., 6379 | WIS | 276 892 | 1 272 | 0.5 | 5 | 1 |
| *Solanum commersonii Poir.* | Boldrini I. & Boechat S., 264 | WIS | 24 6872 | 4 759 | 1.9 | 61 | 58 |
| *Solanum commersonii Poir.* | CST, s.n. | F | 1 057 490 | 11 481 | 1.1 | 69 | 65 |
| *Solanum commersonii Poir.* | Hawkes J.G. et al., 706 | WIS | 162 288 | 2 723 | 1.7 | 33 | 23 |
| *Solanum commersonii Poir.* | Okada K.A., 4583 | WIS | 438 364 | 2 566 | 0.6 | 30 | 20 |
| *Solanum commersonii Poir.* | Okada K.A., 5059 | WIS | 406 536 | 12 707 | 3.1 | 67 | 63 |
| *Solanum malmeanum* Bitter | Brücher E.H., 10 | WIS | 832 765 | 9 272 | 1.1 | 67 | 64 |
| *Solanum malmeanum* Bitter | Brücher E.H., 16 | WIS | 702 991 | 10 665 | 1.5 | 63 | 59 |
| *Solanum malmeanum* Bitter | Brücher E.H., 18 | WIS | 312 294 | 5 938 | 1.9 | 63 | 58 |
| *Solanum malmeanum* Bitter | Brücher E.H., 19 | WIS | 307 622 | 4 119 | 1.3 | 36 | 31 |
| *Solanum malmeanum* Bitter | Delhey R., s.n. | WIS | 1 083 016 | 21 979 | 2 | 74 | 69 |
| *Solanum malmeanum* Bitter | Hawkes J.G. et al., 725 | WIS | 189 964 | 2 257 | 1.2 | 29 | 17 |
| *Solanum malmeanum* Bitter | Krapovickas A. & Cristóbal C.L., 14792 | WIS | 322 990 | 5 209 | 1.6 | 46 | 37 |
| *Solanum malmeanum* Bitter | Okada K.A., 5074 | WIS | 1 027 545 | 23 975 | 2.3 | 71 | 69 |
| *Solanum malmeanum* Bitter | Okada K.A., 7254 | WIS | 591 762 | 12 962 | 2.2 | 67 | 65 |
| *Solanum malmeanum* Bitter | Okada K.A., 7303 | WIS | 523 353 | 11 034 | 2.1 | 69 | 65 |
| *Solanum malmeanum* Bitter | Okada K.A., 7313 | WIS | 137 904 | 4 994 | 3.6 | 51 | 48 |
| *Solanum stoloniferum* Schltdl. | Waterfall U.T. & Wallis C.S., 13489 | BRIT | 164 878 | 4 395 | 2.7 | 52 | 48 |

Values are given per accession, with collector and herbarium, total reads, chloroplast reads mapped, percentage of chloroplast reads relative to total, annotated chloroplast genes recovered, and genes recovered at 75% or more of their length.

For the plastid phylogeny, the *S. chacoense* accession Tressens 4742 recovered no loci and was removed from the analysis.

**Table S3.** Summary of IQ-TREE alignment statistics for nuclear (A353 exons and supercontigs) and chloroplast (exons and supercontigs) sequences.

| **Alignment type** | **Alignment length** | **Number of constant sites** | **Number of variable sites** | **Number of parsimony-informative sites** | **Number of singleton sites** | **Proportion of gaps/missing data** | **Mean GC content** | **Mean entropy per site** | **Informativeness score (Townsend 2007)** |
| --- | --- | --- | --- | --- | --- | --- | --- | --- | --- |
| Chloroplast exons | 63 562 | 61 911 | 1 651 | 39 | 1 612 | 0.26 | 38.54 | 0.15 | 0.74 |
| Chloroplast supercontigs | 105 014 | 96 342 | 8 672 | 710 | 7 962 | 0.40 | 36.68 | 0.03 | 0.71 |
| Nuclear (A353) exons | 209 286 | 197 636 | 11 650 | 345 | 11 305 | 0.17 | 43.24 | 0.17 | 0.56 |
| Nuclear (A353) supercontigs | 943 642 | 797 276 | 146 366 | 11 219 | 13 5147 | 0.42 | 37.45 | 0.11 | 0.69 |

Statistics for the chloroplast and nuclear (exon and supercontig) datasets, including alignment length, constant, variable, parsimony-informative, and singleton sites, proportion of gaps or missing data, mean GC content, mean entropy per site, and the informativeness score **[145]**.

**Table S4.** Mean values and ranges (minimum–maximum) of continuous morphometric traits evaluated in *Solanum chacoense* (northern and southern populations), *S. commersonii*, *S. malmeanum*, and hybrid (*S. commersonii* × *S. malmeanum*).

| **Trait** | ***S. chacoense* N** | ***S. chacoense* S** | ***S. commersonii*** | ***S. malmeanum*** | ***S. commersonii* × *S. malmeanum*** |
| --- | --- | --- | --- | --- | --- |
| **Leaf length (cm)** | 27.14 (19.62–32.84) | 14.67 (9.51–25.88) | 15.91 (6.75–22.08) | 16.20 (10.17–24.67) | 15.53 (12.01–19.05) |
| **Leaf width (cm)** | 16.36 (10.02–20.07) | 8.62 (4.81–12.41) | 6.83 (4.63–9.02) | 6.47 (5.10–9.74) | 5.54 (4.49–6.60) |
| **Leaf length/width ratio** | 1.70 (1.50–2.00) | 1.72 (1.30–2.30) | 2.31 (1.50–3.10) | 2.50 (1.70–2.90) | 2.80 (2.70–2.90) |
| **Length from widest point of leaf to apex (cm)** | 10.89 (5.06–15.29) | 6.35 (3.84–8.91) | 5.04 (2.95–7.18) | 6.48 (3.98–9.24) | 4.55 (4.14–4.96) |
| **Length of terminal leaflet lamina (cm)** | 7.86 (6.34–10.45) | 4.57 (3.00–6.26) | 4.72 (2.44–7.12) | 4.35 (3.55–6.01) | 4.59 (3.82–5.36) |
| **Width of terminal leaflet lamina (cm)** | 3.95 (2.47–5.78) | 2.49 (1.73–3.87) | 2.51 (1.64–3.50) | 2.60 (2.25–2.99) | 1.84 (1.50–2.19) |
| **Length from widest point of terminal leaflet lamina to apex (cm)** | 4.30 (2.91–6.88) | 2.01 (0.95–2.91) | 2.03 (1.33–2.83) | 2.13 (1.60–3.15) | 2.58 (2.25–2.90) |
| **Acumen length of terminal leaflet lamina (mm)** | 3.4 (1.6–5.0) | 2.7 (1.3–4.7) | 1.1 (0–2.0) | 0.8 (0–1.8) | 0.05 (0.00–0.10) |
| **Petiole length of terminal leaflet (cm)** | 1.86 (1.38–2.64) | 1.03 (0.57–1.50) | 0.72 (0.36–0.91) | 0.92 (0.34–1.56) | 0.61 (0.52–0.70) |
| **Distance between lateral leaflets (cm)** | 3.34 (2.74–3.92) | 1.79 (0.84–3.15) | 1.74 (0.58–2.45) | 1.84 (1.27–2.60) | 1.56 (1.29–1.83) |
| **Width of primary lateral leaflet lamina (cm)** | 0.81 (0.69–1.01) | 1.15 (0.62–2.20) | 0.86 (0.60–1.25) | 0.74 (0.39–1.12) | 0.53 (0.51–0.55) |
| **Petiole length of primary lateral leaflet (cm)** | 0.6 (0–0.9) | 1.8 (0.3–2.9) | 1.3 (0.3–2.4) | 0.7 (0.4–1.4) | 0.11 (0.00–0.22) |
| **Width of most distal lateral leaflet lamina (cm)** | 3.50 (2.45–5.08) | 1.88 (1.17–2.85) | 1.54 (1.10–1.94) | 1.58 (1.24–2.36) | 1.20 (0.93–1.47) |
| **Length from widest point of most distal lateral leaflet lamina to apex (cm)** | 3.74 (2.42–6.08) | 1.77 (0.78–2.65) | 1.83 (1.30–2.18) | 1.67 (0.89–2.56) | 1.59 (1.07–2.11) |
| **Petiole length of most distal lateral leaflet (cm)** | 0.42 (0.30–0.49) | 0.27 (0.00–0.51) | — | 0.05 (0.00–0.33) | 0.15 (0.11–0.20) |
| **Peduncle length (cm)** | — | 9.45 (1.97–19.24) | 4.33 (1.68–7.14) | 6.60 (3.71–10.11) | 7.46 (4.50–10.41) |

**Figures**

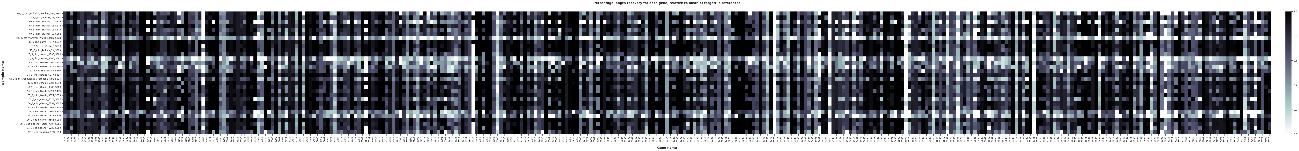

**Supplementary Fig. 1.** Recovery heatmap nuclear.

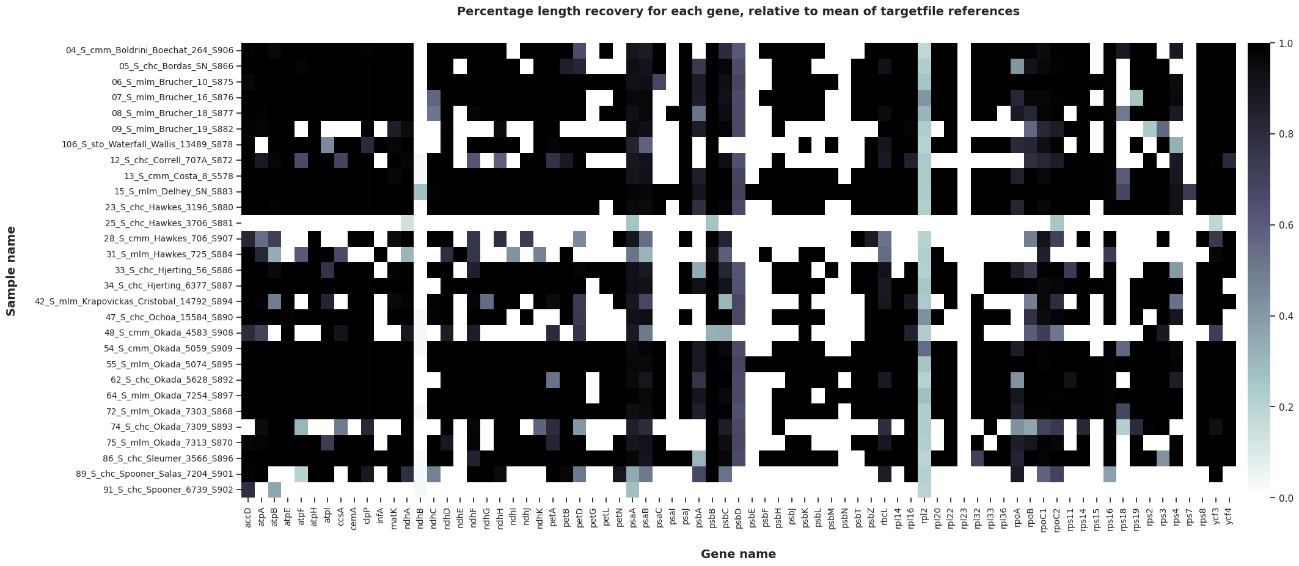

**Supplementary Fig. 2.** Recovery heatmap plastid.

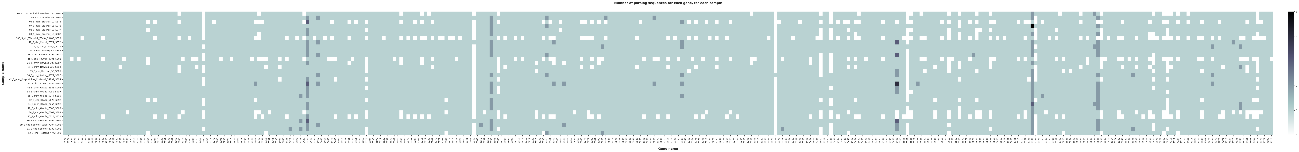

**Supplementary Fig. 3.** Paralog heatmap nuclear.

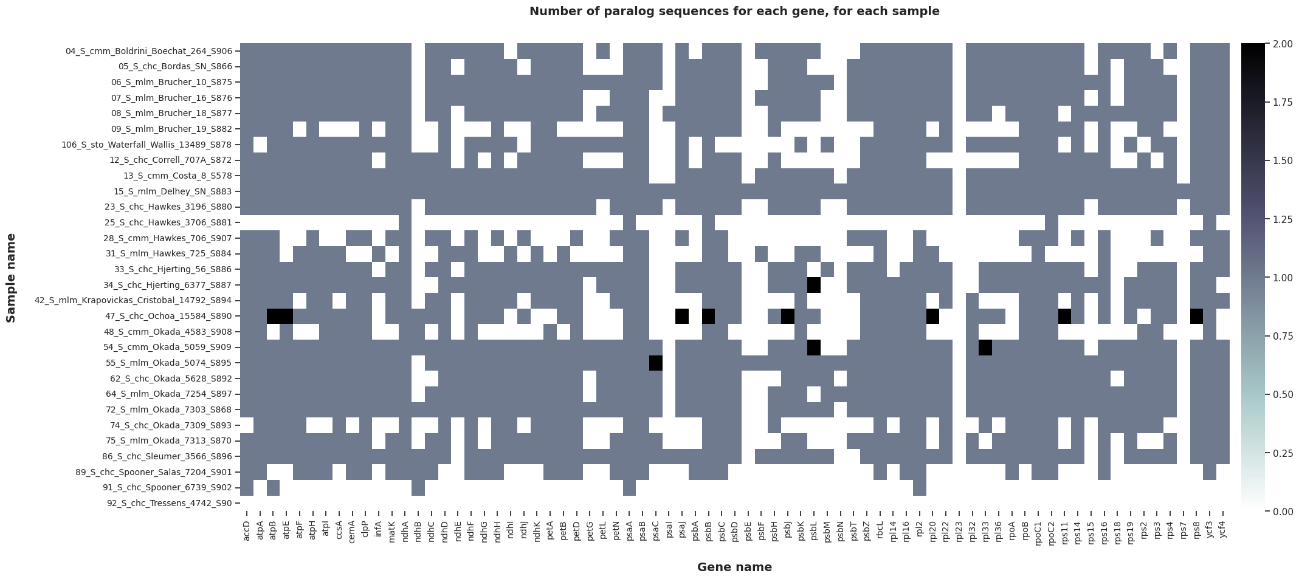

**Supplementary Fig. 4.** Paralog heatmap plastid.

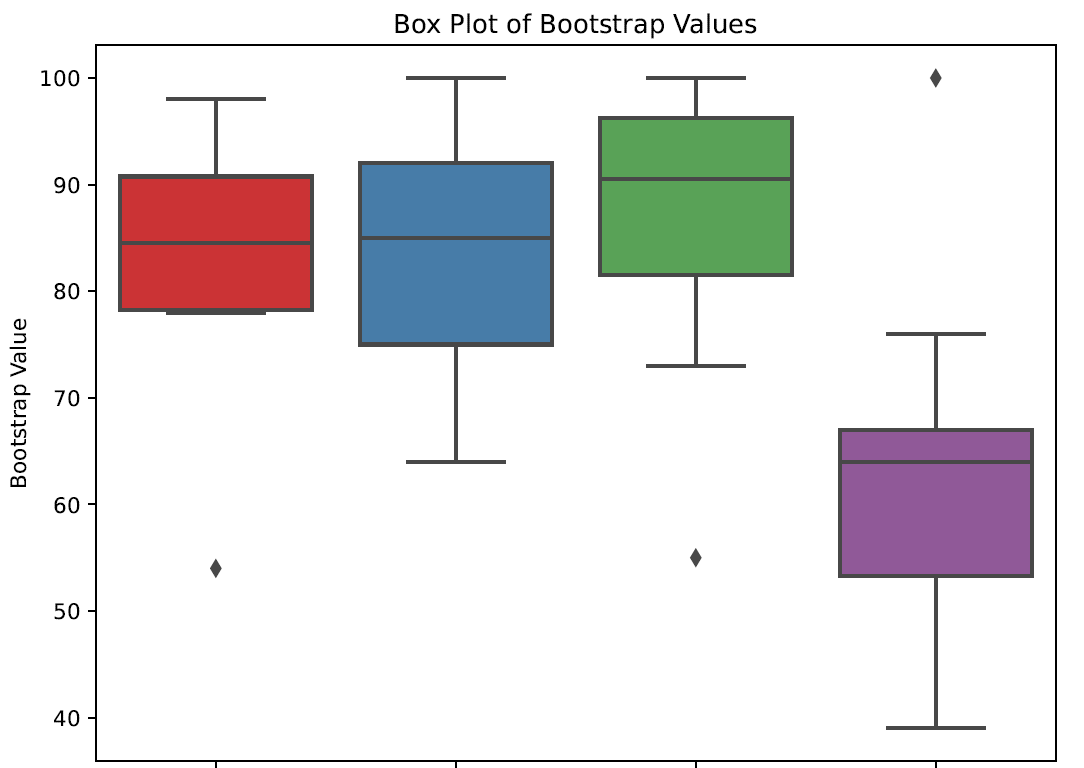

**Supplementary Fig. 5.** Box plots comparing bootstrap values for nuclear (red, supercontigs; blue, exons) and chloroplast (green, supercontigs; purple, exons) sequences.

We carried the supercontigs forward for all phylogenomic analyses because they yielded more variable sites.

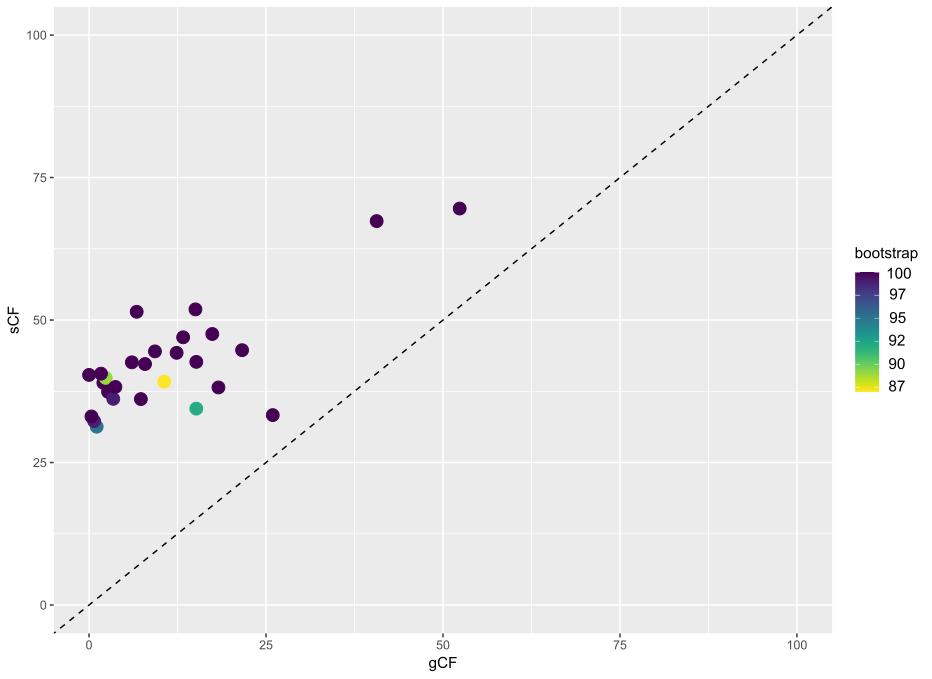

**Supplementary Fig. 6.** Relationship between gene concordance factor (gCF) and site concordance factor (sCF) for the internal branches of the inferred species tree. Each point represents an internal branch, with point color indicating ultrafast bootstrap support (UFBoot). The dashed diagonal line (1:1) represents equal values of gCF and sCF. Most branches exhibit substantially higher sCF than gCF, indicating that while individual gene trees show considerable discordance, the underlying site patterns provide stronger support for the inferred relationships. Most branches received maximal or near-maximal bootstrap support (87–100%), suggesting that the observed gene tree discordance is unlikely to reflect uncertainty in the species tree topology.

1. Blomberg, S.P., Garland, T., Jr., & Ives, A.R. 2003. Testing for phylogenetic signal in comparative data: behavioral traits are more labile. *Evolution* 57(4): 717–745. [https://doi.org/10.1111/j.0014- 3820.2003.tb00285.x](https://doi.org/10.1111/j.0014-%203820.2003.tb00285.x)
2. Revell, L.J. 2012. phytools: An R package for phylogenetic comparative biology (and other things). Meth. Ecol. Evol. 3: 217–223. <https://doi.org/10.1111/j.2041-210X.2011.00169.x>
3. Interpretation of Blomberg’s K for phylogenetic signal: Weak (K > 0.6, not significantly different from 0, not significantly different from 1); Moderate (K < 1, significantly different from 0, not significantly different from 1); Strong (K > 1, significantly different from 0, not significantly different from 1); No phylogenetic signal (K < 0.6, not significantly different from 0, significantly different from 1).
4. Blomberg, S.P., Garland, T., Jr., & Ives, A.R. 2003. Testing for phylogenetic signal in comparative data: behavioral traits are more labile. *Evolution* 57(4): 717–745. [https://doi.org/10.1111/j.0014- 3820.2003.tb00285.x](https://doi.org/10.1111/j.0014-%203820.2003.tb00285.x)
5. Posit team (2025). RStudio: Integrated Development Environment for R. Posit Software, PBC, Boston, MA. URL <http://www.posit.co/>
